# A spatiotemporal single-cell atlas reveals coordinated immune, metabolic, and nutrient exchange programs and a coumarin-centered metabolic switch during soybean arbuscular mycorrhizal symbiosis

**DOI:** 10.64898/2026.07.07.737096

**Authors:** Leonidas D’Agostino, Kaushik Ghose, Yong-Villalobos Lenin, Luis Herrera-Estrella, Gunvant B Patil

## Abstract

Arbuscular mycorrhizal fungi (AMF) establish intimate symbiosis with plant roots, yet the cell-type-specific regulatory and metabolic programs governing this interaction remain poorly resolved. Here, we integrate single-nucleus RNA sequencing (snRNA-seq) with spatial metabolomics across a temporal gradient of soybean root colonization (2-8 weeks post inoculation) to construct a high-resolution, multi-omic atlas of AMF symbiosis. Profiling 33,410 nuclei spanning all major root cell types, we uncover dynamic, cell-type-resolved transcriptional reprogramming coupled to spatially localized metabolite accumulation. Early colonization triggers a robust, epidermis-localized immune response alongside cortex-specific epigenetic reprogramming, mediated by RNA-directed DNA methylation machinery, suggesting active suppression of defense in fungal accommodation zones. Spatial metabolomics reveals a biphasic metabolic transition from flavonoid- and terpenoid-rich signaling states to lipid-dominated nutrient exchange, aligned with colonization progression. In parallel, coordinated carbon allocation and lipid biosynthesis pathways were activated in cortex and vascular tissues, supporting fungal dependence on host-derived fatty acids and sugars. Nutrient exchange programs, particularly nitrogen and phosphorus transport, exhibit strong pericycle and phloem specificity, highlighting systemic integration of symbiotic benefits. Through co-expression network analysis, we identify a previously uncharacterized coumarin-centered metabolic switch, governed by *GmF6’H1-2*, that is essential for efficient colonization, as validated by natural loss-of-function variants. Collectively, this study provides a comprehensive, spatially resolved framework linking gene regulation, metabolism, and cell identity, revealing that AMF symbiosis is orchestrated through coordinated immune modulation, metabolic rewiring, and nutrient flux partitioning at single-cell resolution.

**Key points:**

- Multi-omic integration reveals cell-type–resolved symbiotic programs
- Symbiosis requires spatially coordinated immune reprogramming
- A biphasic metabolic shift underpins colonization dynamics
- Carbon and nutrient flux are partitioned across specialized cell types
- Discovery of a coumarin-driven “metabolic GO-switch” controlling symbiosis

## Introduction

Arbuscular Mycorrhizal Fungi (AMF) form one of the most ancient and ecologically consequential symbioses on Earth, colonizing the roots of over 80% of land plant species [1, 2]. Unlike ectomycorrhizal fungi, which remain confined to the root surface and intercellular spaces via the Hartig net [3], AMF penetrate the root epidermis and differentiate specialized intracellular structures (arbuscules) within cortical cells. These branched hyphal trees dramatically expand the plant-fungus interface, enabling efficient bidirectional nutrient exchange that is fundamental to plant fitness. Through this interface, AMF delivers soil-derived phosphorus (P), nitrogen (N), and water directly to the host, while the plant reciprocates by allocating up to 20% of its photosynthetically fixed carbon, predominantly as fatty acids and sugars, to sustain fungal growth and metabolism [4–6].In general, the molecular dialogue facilitating AMF symbiosis is well-established, wherein the host roots release strigolactones and flavonoids into the rhizosphere, stimulating fungal branching and metabolic activation. AMF reciprocate with lipochitooligosaccharide and chitooligosaccharide signals that activate the conserved symbiosis signaling pathway (CSSP) in the host [7]. In the case of soybeans and other legumes, the CSSP is shared with nitrogen fixing rhizobia bacteria, yet AMF symbiosis is distinguished by its broad host range and lack of strict host specificity, though ecological host preferences and niche partitioning have been documented [8, 9]. Despite these foundational insights, the precise cellular and molecular events governing fungal recognition, epidermal penetration, cortical accommodation, and ultimately arbuscule development and senescence remain incompletely understood. A central challenge is that AMF colonization is inherently asynchronous. At any given stage, a single root simultaneously harbors fungal structures at vastly different developmental stages, from initial hyphopodia at the epidermis to fully mature arbuscules and collapsing senescent structures deep in the cortex [10]. This biological heterogeneity necessarily confounds conventional whole-root transcriptomic approaches, which average signals across millions of cells in different symbiotic states, thus masking the cell-type-specific and stage-specific regulatory programs that drive each phase of colonization [11].

Although laser-capture microdissection (LCM) has been employed to obtain targeted transcriptomes of cortex cells at known colonization stages, it is inherently limited by the requirement for prior anatomical knowledge. Additionally, it captures limited cells, and cannot simultaneously resolve the full complement of root cell types engaged in symbiosis [12]. The emergence of single-nucleus RNA sequencing (snRNA-seq) now offers a transformative alternative through unbiased, high-resolution profiling of gene expression across all cell types within tissues, capturing continuous transcriptional states without prior assumptions about cell identity or colonization stage [13–15]. In parallel, spatial metabolomics has emerged as a powerful complement to transcriptomics, enabling the in-situ mapping of thousands of metabolites at near-cellular resolution within tissue sections, providing direct readout of the biochemical outputs of gene regulatory programs [16, 17]. Together, these technologies create an unparalleled opportunity to link cell identity, transcriptional regulation, and metabolic state within the spatial context of the colonized root.

While early single-cell transcriptomic efforts in model legumes and other plants have begun to reveal cellular heterogeneity during AMF colonization [13, 18], the dynamic metabolic rewiring accompanying transcriptional reprogramming remains uncharacterized, and a comprehensive spatiotemporal atlas of AMF symbiosis in a major crop species is entirely lacking. Soybean, the world’s most economically important legume crop, presents a uniquely compelling system. It’s shared symbiotic signaling pathway with nitrogen-fixing rhizobia introduces regulatory complexity not present in non-legume hosts and its agronomic importance makes mechanistic insights directly translatable to sustainable agriculture. To address these gaps, we constructed a high-resolution, spatiotemporal multi-omic atlas of AMF symbiosis in soybean by profiling 33,410 nuclei via snRNA-seq across a temporal gradient spanning 2, 4, 6, and 8 weeks post-inoculation, capturing the full arc from initial colonization to established symbiosis alongside an uninoculated control. Complementing this transcriptional atlas, spatial metabolomics was applied to two early colonization time points (2 and 4 weeks) to map metabolite distributions at near-cellular resolution within intact root sections. Critically, key regulatory and metabolic genes emerging from this atlas are functionally validated through natural loss-of-function variants identified in whole-genome resequencing germplasm collections, directly linking molecular discovery to biological function. This integrated framework reveals dynamic, cell-type-resolved transcriptional reprogramming coupled to spatially explicit metabolic transitions, uncovers a previously uncharacterized coumarin-mediated metabolic checkpoint essential for colonization efficiency, and establishes a comprehensive molecular foundation for understanding and ultimately engineering AMF symbiosis in a major crop species.

## Results

### snRNA-seq atlas of soybean root AMF symbiosis

To understand cell-type specific transcriptional profile of plant root-AMF interaction, we performed snRNAseq on soybean roots sampled across a temporal colonization gradient spanning 2, 4, 6 and 8 weeks post inoculation (wpi), alongside an uninoculated control harvested at the 6-week time point, collectively capturing the progressive stages of fungal establishment, colonization expansion, and mature symbiosis (Fig. 1A).

**Fig. 1.**
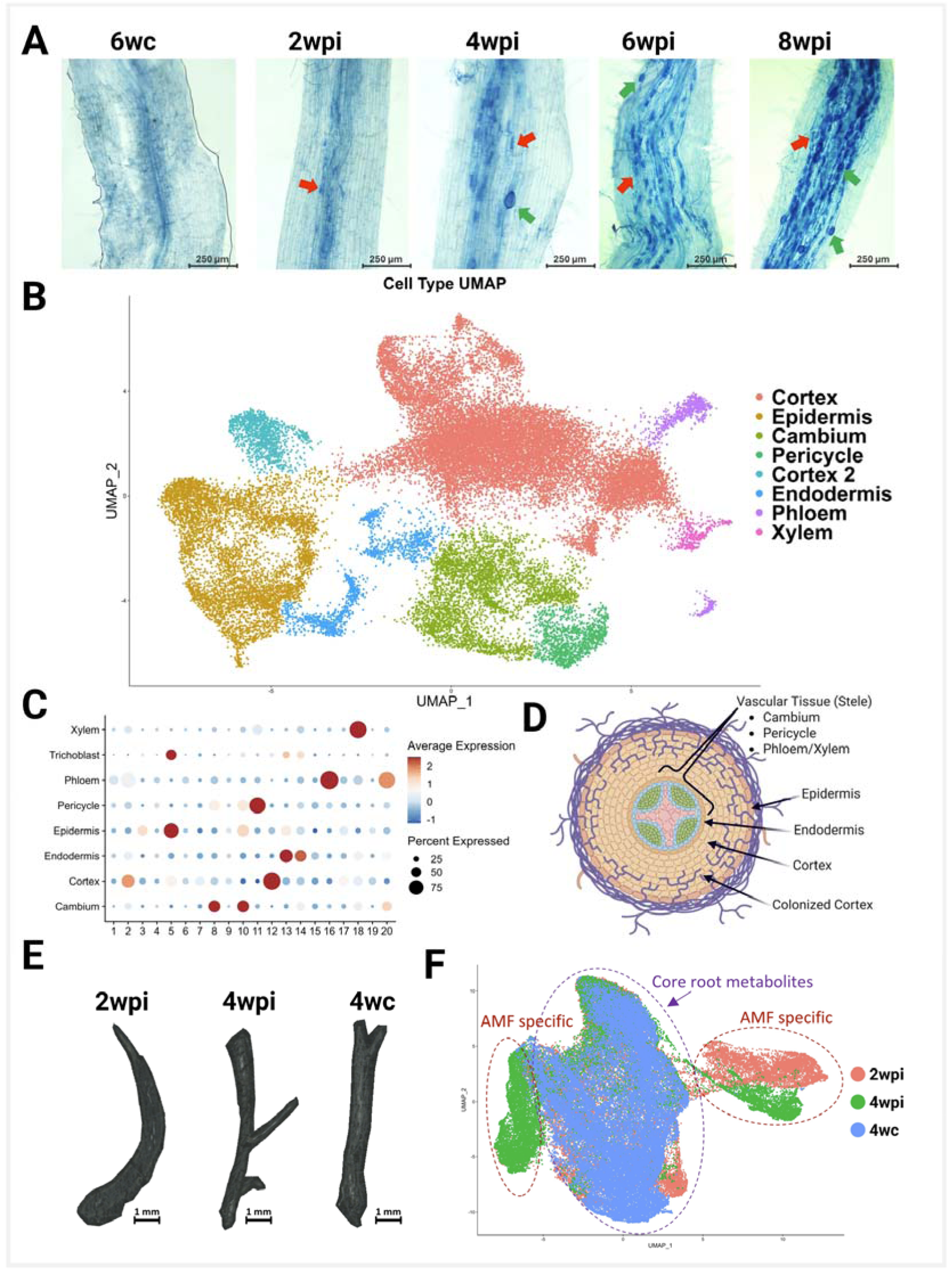
Single-nucleus RNA sequencing and spatial metabolomics atlas of soybean root AMF symbiosis. **(A)** Trypan blue-stained soybean root sections from uninoculated control (6wc) and AMF-inoculated roots at 2, 4, 6, and 8 weeks post inoculation (wpi). Red arrows indicate arbuscules; green arrows mark vesicle formation in cortical cells. Scale bars = 250 µm. **(B)** Integrated UMAP projection of 33,410 high-quality nuclei across all five timepoints (2, 4, 6, 8wpi and 6wc), resolved into 20 transcriptionally distinct clusters and annotated to eight major root cell-type identities. Each color represents a cell type as indicated in the legend. **(C)** Consensus dot plot of marker gene expression across all 20 clusters confirming cell-type annotation specificity. Dot size indicates the percentage of nuclei expressing each marker gene set; color intensity represents scaled average expression (blue = low, red = high). **(D)** Schematic cross-section of the soybean root illustrating the radial organization of annotated cell types from the epidermis to the vascular stele, with the cortex depicted as the primary zone of hyphal network establishment and arbuscule formation. Made in Biorender. **(E)** Representative soybean root segments used for untargeted spatial metabolomics at 2wpi, 4wpi, and 4wc, showing the morphological diversity of colonized versus uninoculated roots. Scale bars = 1 mm. All root segments are present in Fig. S2 **(F)** UMAP projection of spatial metabolomics data in which each pixel (50 µm²) is treated as an independent metabolite unit and dimensionality reduction was computed across the full detected metabolome. Each color represents a sample as indicated in the legend.

Following quality control and filtering, 33,410 high-quality nuclei were retained for downstream analysis, 7,263 (2wpi), 6,795 (4wpi), 7,522 (6wpi), 7,993 (8wpi), and 7,010 (6wc), with a mean of 517 unique molecular identifiers (UMIs) and 601 expressed genes per nucleus (Table S1). Collectively, these nuclei captured 45,743 soybean genes, representing 86.52% of the 52,872 annotated genes in the Glycine max Wm82 v4 reference genome, demonstrating broad transcriptomic coverage across the atlas (Fig. 1b). Integrated analysis of all five samples in Seurat [19] resolved 20 transcriptionally distinct clusters (Fig. S1A), which were annotated to major root cell types using a curated set of known and newly validated soybean marker genes [20–23] (Table S2). This annotation assigned clusters to all principal root tissue layers: xylem (cluster 18), phloem (clusters 16 and 20), cambium (clusters 8 and 9), pericycle (cluster 11), endodermis (clusters 13 and 14), epidermis (clusters 3, 5, and 19), and cortex (clusters 1, 2, 4, 6, 7, 9, 15, and 17) (Fig. 1B,C, S1B). Marker gene expression across all clusters was visualized in a consensus dot plot (Fig. 1c), confirming the specificity and robustness of cell-type annotations. Notably, cluster 12 exhibited disproportionately high expression of cortex marker genes relative to all other clusters while occupying a distal position in the UMAP embedding (Fig. S1B), indicating a transcriptionally specialized cortex state (hereafter designated cortex 2). The resulting UMAP projection (Fig. 1B) and schematic tissue illustration (Fig. 1D) together provide a spatially and transcriptionally coherent reference framework for all subsequent cell-type-resolved analyses of AMF symbiosis.

### Spatial metabolomics reveals a biphasic metabolic transition during AMF colonization

To map the spatial and temporal metabolic landscape of AMF symbiosis at near-cellular resolution, we performed spatial metabolomics on soybean root sections from 2wpi and 4wpi AMF-inoculated samples alongside a 4-week uninoculated control (4wc) (Fig. 1E, S2). Each pixel (50 µm^2^) was treated as an independent metabolite unit, and dimensionality reduction via UMAP was computed across the full detected metabolome using the Seurat package in R [19] (Fig. 1F). As evidenced, 4wc pixels overlapped broadly with both AMF-treated time points, yet 2wpi and 4wpi pixels diverged substantially from the control showing AMF specific clusters in distinct directions (Fig. 1F). Unsupervised clustering of all pixels distinguished eight metabolic clusters (Fig. 2A). When samples were examined individually, clusters 4 and 8 emerged as AMF-specific (absent from the 4wc control) with cluster 4 predominating at 2wpi and cluster 8 at 4wpi (Fig. 2B). Spatial projection of these clusters onto root tissue sections revealed a distinct temporal partitioning, wherein cluster 4 localized predominantly to root segments 2wpi-i, -vii, -ix, and 4wpi-iv, while cluster 8 defined segments 4wpi-i, -ii, -iii, and vi to ix (Fig. 2c). Hereafter we designated these as 2wpi-colonized and 4wpi-colonized states, respectively, with segments lacking either cluster classified as non-colonized.

**Fig. 2.**
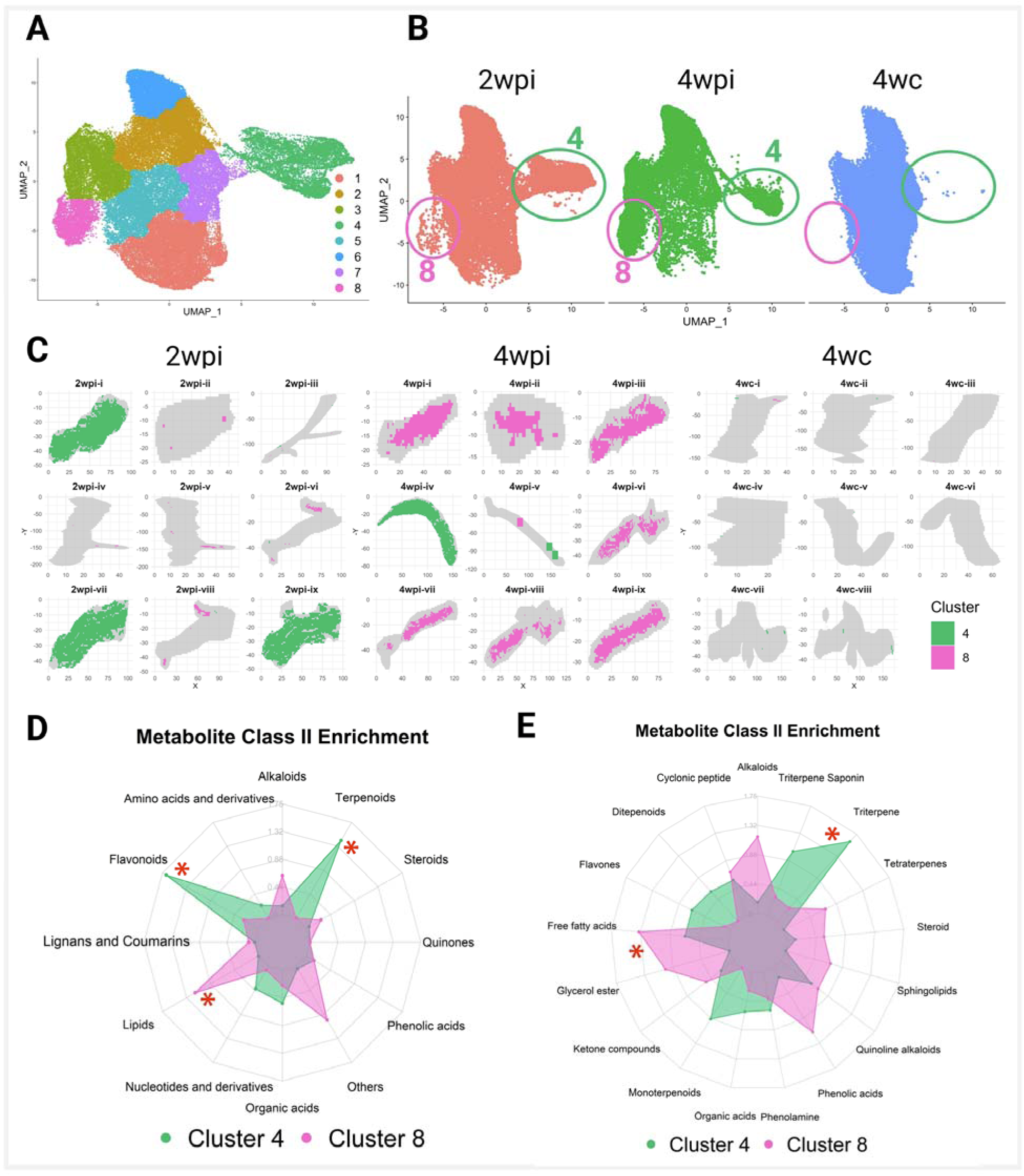
Spatial metabolomics reveals AMF-specific metabolic states and a temporal shift from signaling to lipid-dominated symbiosis. **(A)** Unsupervised clustering of all spatial metabolomics pixels resolved eight distinct metabolic clusters. **(B)** Re-clustering of individual samples showed cluster 4 and 8 specific to AMF treatment (2 and 4 wpi) compared to the 4-week control (4wc). **(C)** Spatial mapping of these clusters onto root sections delineated distinct colonization states. Cluster 4 metabolites (green) predominantly localized to 2wpi and few 4wpi samples, while cluster 8 metabolites (pink) occupied predominantly to 4wpi samples. Segments lacking either cluster were designated as non-colonized. **(D–E)** Metabolite class enrichment analysis (Class I: broad categories; Class II: subclasses) revealed distinct biochemical profiles for the two AMF-associated clusters. Cluster 4 (2wpi-colonized) showed significant enrichment of flavonoids and terpenoids (p < 0.05), with triterpenes as the significantly enriched subclass (p < 0.05). Cluster 8 (4wpi-colonized) was characterized by significant enrichment of lipids (p < 0.05), with free fatty acids as the significantly enriched subclass (p < 0.05). Red asterisks denote statistically significant enrichment (p < 0.05).

Metabolite class enrichment analysis (Class I: broad biochemical categories; Class II: specific subclasses) of clusters 4 and 8 revealed a biologically coherent metabolic transition (Fig. 2D,E). Cluster 4 (2wpi-colonized) was significantly enriched in flavonoids and terpenoids (p < 0.05), with triterpenes representing the most enriched subclass, which is reflective of the known roles of flavonoids as pre-symbiotic host signals that stimulate hyphal branching and of terpenoids as modulators of early immune response [24, 25]. Organic acids and amino acids showed non-significant enrichment trends, and free fatty acids were moderately elevated, potentially reflecting the onset of carbon reallocation toward the developing fungal interface (Fig. 2D). In contrast, cluster 8 (4wpi-Colonized) was dominated by a significant enrichment of lipids, particularly free fatty acids (p < 0.05), alongside phenolic acids and alkaloids, while flavonoid enrichment was markedly reduced relative to cluster 4 (Fig. 2E). This metabolic shift from a signaling-competent state toward a lipid-dominated biochemical environment is mechanistically consistent with the established dependence of AMF on host-derived fatty acids as a primary carbon source during active symbiosis [26]. The spatial resolution of this transition across individual root segments provides a metabolic framework that complements and contextualizes the transcriptional atlas described below.

### Defense modulation exhibits a cell-type specific pattern

While AMF are beneficial microbes, a fundamental paradox of AMF symbiosis is that the host plant must simultaneously suppress the immune response and activate symbiosis function [27]. How this immune balance is achieved at the level of individual root cell types has remained poorly understood. Our snRNA-seq atlas reveals that defense pathway activation during AMF colonization is not uniform across root tissues but is instead organized into a precise, cell-type-resolved pattern across colonization time points (Fig. 3). At 2wpi, the epidermis, the first cell layer encountered by AMF, exhibits the most pronounced defense activation, with significant enrichment of pathways involved in jasmonic acid (JA), salicylic acid (SA), response to pathogens, and cell wall–associated defense mechanisms (Fig. 3a). This epidermal defense response was progressively downregulated at 4wpi and 6wpi. This pattern is largely similar in pericycle and cambium cells, while the cortex, which is the primary site of arbuscule formation, shows a near absence of these canonical defense pathways (Fig. 3a). Only the GO terms *defense response to other organism*, *DNA Methylation*, and *regulation of immune response* showed differential detection in cortex and cortex 2 cells at 2wpi (Fig. 3a, S3). This cortex-specific transcriptional defense modulation suggests that the cortex does not passively tolerate fungal entry but instead fine-tunes it to permit arbuscule accommodation while retaining a basal level of pathogen surveillance. This interpretation is supported by the absence of NBS-LRR gene expression in the cortex, with diminishing expression as symbiosis progresses (Fig. S4a). Spatial metabolomics corroborated and spatially resolved this defense architecture, revealing three biochemically distinct responses occurring synchronously within colonized root segments (Fig. 3b). In the outer regions (encompassing the epidermis and outer cortex) of 2wpi-colonized roots, coumarins were the dominant metabolite class, consistent with their known roles as antimicrobial compounds and iron mobilization at the plant-microbe interface [28]. Simultaneously, the inner and middle regions (encompassing the vasculature and inner cortical cells) of 2wpi-colonized segments were characterized by the presence of defense related proanthocyanidins, triterpenes, terpenes, triterpene saponins, and steroidal saponins (Fig. 3b).

**Fig. 3.**
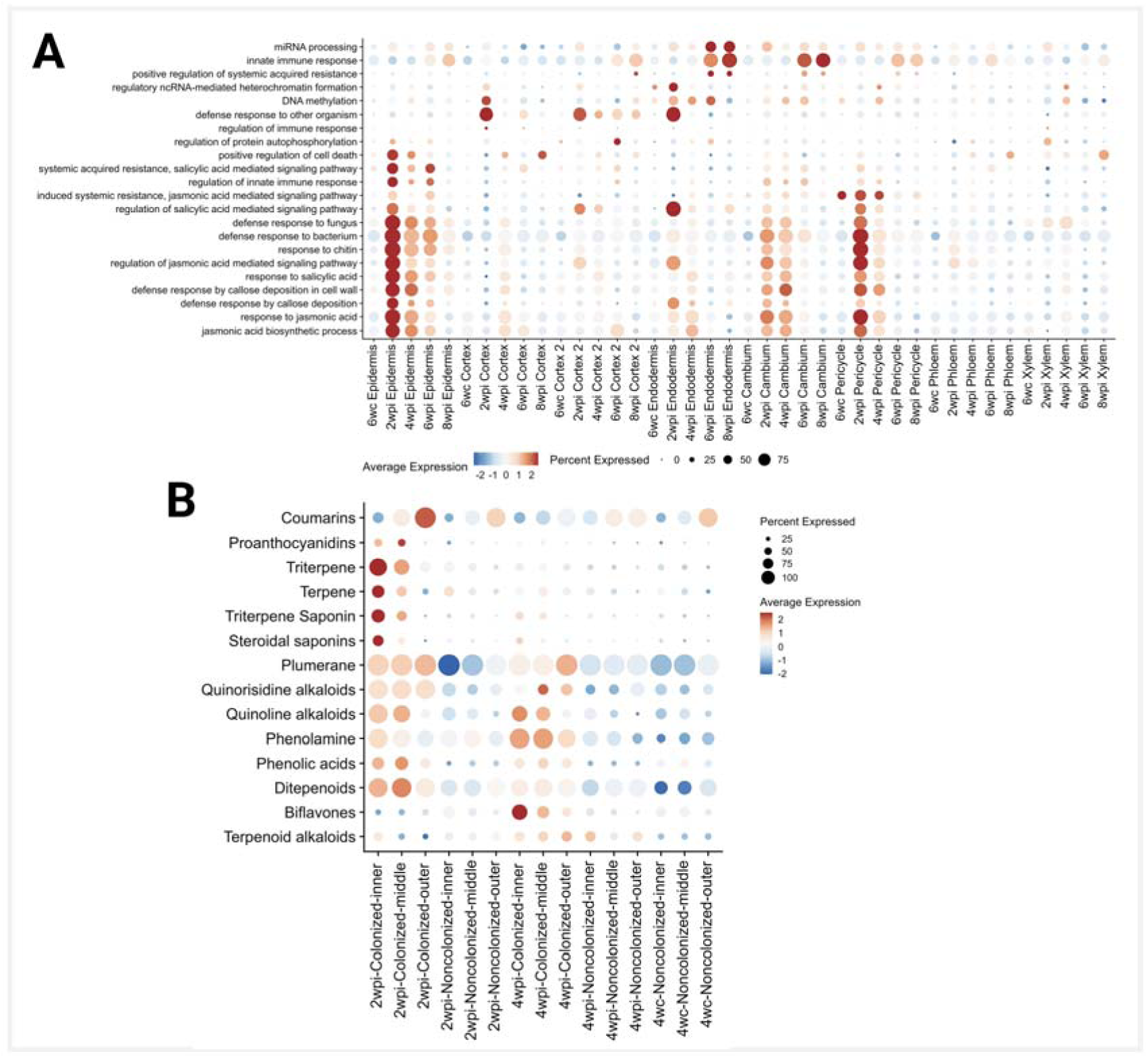
Defense modulation and priming mechanisms in response to AMF. **(A)** Consensus dot plot of defense pathways gene expression across all cell-type sample groups. Dot size indicates the percentage of nuclei expressing each pathway; color intensity represents scaled average expression (blue = low, red = high). Specific pathways are further elucidated in Fig. S3 and S4. **(B)** Consensus dot plot of defense metabolite subclasses across all sample, colonized segment, and region comparison groups. Dot size indicates the percentage of pixels expressing each subclass; color intensity represents scaled average expression (blue = low, red = high).

AMF is also known to promote plant defenses against pathogenic fungi and bacteria, often termed induced systemic resistance (ISR) or mycorrhizal induced resistance (MIR) [29]. Genetically, we see the expression of genes involved in *induced systemic resistance, Jasmonic acid mediated signaling pathway* upregulated in a similar manner to the other defense genes, but expression was also elevated in 6wc pericycle cells (Fig. 3a, S4b). Accumulating synergistically are several defense related metabolites showing near-constitutive detection, including plumeranes, quinorisidine alkaloids, quinoline alkaloids, phenolamine, phenolic acids, and diterpenoids detected at 2wpi, and biflavones and terpenoid alkaloids detected at 4wpi (Fig. 3b). Moreover, gene network analysis revealed the brown gene co-expression network to be primarily localized to the 2wpi and 4wpi epidermal cells, followed by endodermal and cambium cells (Fig. S5, S6). The brown network was significantly enriched in defense related GO terms involved in camalexin and glucosinolate production, SA metabolism and JA signaling, defense, and response to chitin (Fig. S6b). Core hub genes included *GmNSL1, GmPAD4* and *GmJAZ1* (Fig. S6c, table S3).

### Modulation of host carbon partition to support AMF development

Carbon allocation to the fungal symbiont represents one of the most metabolically important commitments that a host plant makes during AMF symbiosis to sustain growth and development [4]. Our integrated snRNA-seq and spatial metabolomics analysis reveals a coordinated, spatially resolved program of sucrose mobilization, fatty acid biosynthesis, and lipid transport that intensifies with colonization progression and is partitioned across functionally distinct root tissues (Fig. 4a). Sucrose transport genes (*GmSUC2-1, GmSUC2-3* and *GmSUC2-6*) were upregulated in phloem cells and pericycle cells of AMF treated roots (Fig. 4A, S7). Downstream of transport, sucrose catabolism genes (*GmSUC4; GmCINV2* and *GmA/N-InvC-2*) involved in sucrose synthase and invertases, were broadly upregulated across multiple cell types at 2wpi, including the epidermis, cortex 2, endodermis, and pericycle, while showing pronounced induction in xylem cells (Fig. 4A, S7B).

**Fig. 4.**
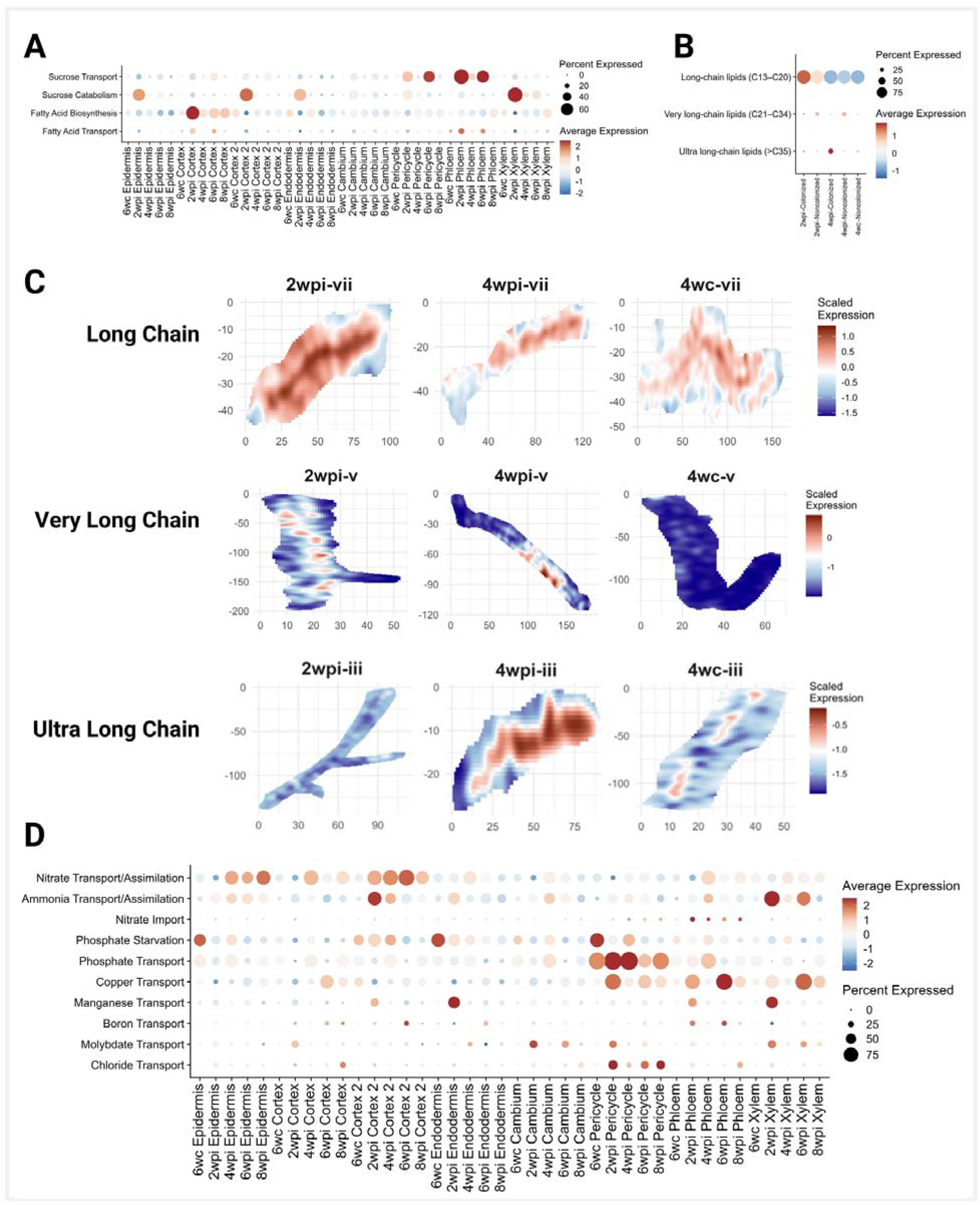
Carbon and nutrient flux changes in relation to AMF presence and plant age. **(A)** Consensus dot plot of carbon pathways gene expression across all cell-type sample groups. Specific pathways are further elucidated in Fig. S7 and S8A. **(B)** Consensus dot plot of lipids grouped by carbon chain length across all sample and colonized segment comparison groups. Dot size indicates the percentage of pixels expressing each group; color intensity represents scaled average expression (blue = low, red = high). **(C)** Representative root plots of data from **(B)**, color intensity represents scaled average expression (blue = low, red = high). Full root plots are present in Fig. S10. **(D)** Consensus dot plot of nutrient transport pathways gene expression across all cell-type sample groups. Dot size indicates the percentage of nuclei expressing each pathway; color intensity represents scaled average expression (blue = low, red = high). Specific pathways are further elucidated in Fig. S11 and S12.

Concurrently, FA biosynthesis was strongly and specifically induced in cortex and endodermal cells across AMF-treated samples, with a particularly pronounced upregulation at 2wpi that was sustained across later time points relative to the uninoculated 6wc control (Fig. 4A, S8A), a pattern consistent with lipid synthesis reprogramming induced my AMF symbiosis localized to the cortex [30]. Analysis of the 50 highest expressed FA genes showed a temporal stage specific shift in biosynthesis programs (Fig. S8A). Compared to control sample (6wc), early colonization (2wpi) cortex cells were enriched in *de novo* FA biosynthesis genes associated with acetyl-CoA carboxylase (CAC), acyl carrier protein (ACP), and enoyl-ACP reductase (MOD1), while later time points (4–8wpi) showed progressive enrichment for genes associated with fatty acid export (FatB), β-ketoacyl-CoA synthase (KCS), and wax biosynthesis (CER) (Fig. S8a). Several core FA biosynthesis genes *GmEMB3147, GmBCCP2, GmCYP77B1-2*, and *GmKASI-2* were consistently elevated across all AMF time points relative to 6wc, indicating constitutive symbiosis-driven induction of specific FA biosynthetic machinery (Fig. S8a).

In concordance with the snRNAseq analysis, spatial metabolomics reveals that long-chain lipids (C chain length 13-20) were highly upregulated in 2wpi root segments, with a higher enrichment in 2wpi-colonized segments. (Fig. 4b, c, S9). Additionally, very long-chain lipids (C21-C34) showed a specific enrichment in 2wpi and 4wpi noncolonized tissue, reflecting the flux of short chain FA synthesis genes observed in the snRNA-seq (Fig. S8), while ultra long-chain lipids (>C35) showed a discrete upregulation in 4wpi-colonized segments, suggesting a spatial and temporal segregation of lipid species between colonized and non-colonized tissue zones. At the level of individual lipid species, membrane-associated long-chain fatty acid lipids including LysoPC 16:0, LysoPC 14:0 (2n isomer), and PS(14:1/18:4) showed increased abundance in the inner and middle regions of 2wpi root segments, corresponding to putative vascular and cortical zones, while 1-Palmitoyl-2-(5-hydroxy-8-oxo-6-octenedioyl)-sn-glycero-3-phosphatidylcholine peaked specifically in the outer colonized region at 2wpi, potentially marking the lipid transfer interface at the epidermal-cortical boundary (Fig.8B, S10).

To decipher hexose synthesis and mobilization, three glucose-containing saccharides D-maltotetraose (4 glucose containing), acetyl-maltose (2 glucose containing), and maltotriose (3 glucose containing) were mapped on root sections. D-maltotetraose and acetyl-maltose showed differential localization in the colonized 2wpi and 4wpi root segments, while maltotriose was detected more in the colonized 2wpi and 4wpi outer root segments. Lastly, to capture general lipid metabolism, methyl stearate (18:0; stearic acid) and methyl ricinoleate (18:1; oleic acid) were mapped on roots. Both showed high preferential accumulation in colonized root segments, with methyl stearate localized to the outer regions and methyl ricinoleate to the inner and middle regions (Fig. S8b, S10).

### Nutrient flux dynamics influenced by AMF symbiosis

A well-studied function of AMF symbiosis is the improved nutrient uptake of host plant, mainly N and P though the hyphal network [5, 6]. However, coordination of these nutrient uptake programs at the individual cell level and remain difficult to decipher. This study shows that AMF colonization triggers a comprehensive, cell-type resolved reorganization of nutrient transport and assimilation genes expression (Fig. 4D, E). In the case of N dynamics, AMF colonization broadly upregulated genes involved in nitrate and ammonia transport and assimilation across epidermis, cortex, and cortex 2, with higher expression relative to 6wc (Fig. 4D). Nitrate import was highly upregulated in AMF phloem tissue, driven primarily by *GmCEPR1-1* and *GmCEPR1-2* (Fig. 4F, S11A). Several nitrate transport genes, *GmNRT3.1-1/2, GmNRT2.4-1/3/* and *4* show increased expression in 4wpi to 8wpi epidermal cells and 8wpi cortex cells (Fig. S11B). *GmAMT2-5/8* followed a similar pattern and *GmAMT2-1,4* and *GmAMT1;2-1* showed the highest expression at 8wpi (Fig. S11C). Glutamate transport (GLT) genes *GmGLT1-2/3/4* were spatially expressed in xylem cells, and *GmGLT1-1* was confined to cortex 2 cells (Fig. S11C).

The P response also revealed a biologically distinction between P acquisition and P starvation (Fig. 4D, S12). The 6wc uninoculated roots, representing chronic P stress showed upregulation of P-starvation genes in pericycle, epidermis and endodermis cell layers (Fig. 4F). Interestingly, the P starvation response was also slightly active in the 2wpi and 4wpi cortex 2 cells (Fig. 4D, S12A). Several P starvation gene homologs, including PHOSPHATE STARVATION RESPONSE (PHR) genes such as *GmPHR1-1/2/3/* and *4* showed elevated expression in 2wpi and 4wpi cell types over 6wc, namely the pericycle, endodermis, and cortex 2, suggesting AMF symbiosis may activate separate P starvation gene paralogs, but additional experimentation is needed to conclude this. The previously identified *GmPHR25*, a master regulator for P starvation in soybean [31], showed a strong upregulation in 6wc across multiple cell types (Fig. S12A). Likewise, the canonical phosphate-starvation response genes *GmSIZ1*⍰*1*, *GmNPC4*, *GmSPX*, and *GmSQD2* genes [32] were strongly enriched in the epidermis, cortex 2, endodermis, and pericycle of 6wc roots, whereas their expression remained comparatively low in AMF inoculated root cell types (Fig. S12a).

P transport machinery was strongly and specifically induced in pericycle cells across AMF-treated time points, consistent with pericycle-mediated P loading into the vasculature for systemic distribution following fungal delivery at the arbuscular interface [33] (Fig. 4d). Critically, *GmPHO1-1/3/4* and *GmPHO2-4/5* exhibited a clear AMF-associated upregulation in pericycle cells, while *GmPHO1-2* and *GmPHO2-2/3* showed elevated expression specifically in 6wc pericycle cells (Fig. S12b), suggesting a paralog-level functional divergence that distinguishes symbiotic P export from starvation-driven P scavenging, and suggests that AMF colonization selectively recruits a distinct PHO transporter repertoire for symbiosis-compatible P loading. Beyond N and P, AMF colonization induced a broader micronutrient acquisition program in a cell-type–specific manner (Fig. 4d). Copper, molybdate, and chloride transport genes were preferentially upregulated in pericycle cells, while manganese and boron transport genes showed phloem-specific induction indicating a spatial partitioning that reinforces the pericycle as a central symbiotic nutrient integration hub and implicates the phloem in the systemic relay for micronutrients to distal tissues.

### Analysis of cortex cells reveals colonization specific programming

The cortex is the primary site of arbuscule formation and symbiosis, but its transcriptional heterogeneity, spanning cells at vastly different stages of fungal accommodation is entirely masked in whole-root analyses. To resolve this heterogeneity and reconstruct the molecular programs governing each stage of cortical symbiotic development, we performed sub-clustering and trajectory analyses on cortex and cortex 2 cells, enabling identification of colonized cell states and pseudotemporal ordering of their transcriptional trajectories.

UMAP re-clustering of cortex and cortex 2 across all samples showed spatially separate but transcriptionally similar clusters (Fig. 5a). RNA velocity analysis of the combined cortex clusters confirmed the relationship, demonstrating a strong directional splicing trajectory flowing from cortex 2 toward cortex cells (Fig. 5b), suggesting that cortex 2 is a transcriptionally upstream, potentially less differentiated cortical population. Complementing this, co-expression network analysis identified the green gene network as specifically active in 6wpi and 8wpi cortex cells, with secondary activity in phloem (Fig. S7, S13). This network was significantly enriched in GO terms including *copper ion homeostasis*, *water transport*, and *cellular homeostasis*, suggesting that late-stage cortical symbiotic programs extend beyond canonical nutrient exchange to encompass broader ionic and osmotic homeostasis functions (Fig. S13). To identify colonized cortex cells within this population, we analyzed the expression of known AMF marker genes (Table S4) using a module scoring approach [13], designating the top 5% of expressing nuclei as colonized and visualizing their distribution on the cortex UMAP (Fig. 5c). This allowed marker-guided identification of colonized cells without anatomical assumptions, enabling direct comparison of expression between colonized and non-colonized cortex cells. Projecting cortex specific defense and epigenetic regulatory genes onto colonized versus non-colonized cell populations revealed that RdDM pathway genes (*GmCMT3-1/2, GmNRPD2A-1/2, GmNRPD1B-1, GmET2, and GmAGO4-2/4/5*) were specifically induced in colonized cortex cells at 2wpi (Fig. S14a), confirmative that immune suppression is targeted to cells actively undergoing fungal accommodation rather than a broad cortical priming response. In contrast, chaperone genes (*GmHSC70-1/2/5*) [34] were elevated in both colonized and non-colonized 2wpi cortex cells (Fig. S14b), suggesting a broader tissue-level preparation that precedes and is independent of direct fungal contact. Colonized cells at 2wpi additionally showed strong induction of cell cycle–associated pathways consistent with the cytological remodeling and periarbuscular membrane biogenesis demands of arbuscule formation, as well as a documented response to CSSP-induced symbiosis in model organisms (Fig. S14c) [35]. Interestingly, transmembrane trafficking programs followed a temporally ordered transition wherein endocytic pathways dominated in 2wpi-4wpi colonized cells suggesting the shift from arbuscule biogenesis to active nutrient secretion at the periarbuscular interface. Together, these observations delineate a coordination in molecular sequence (epigenetic gating, cellular remodeling, and membrane trafficking), that defines the early-to-mature progression of cortical symbiotic development (Fig. S14d).

**Fig. 5.**
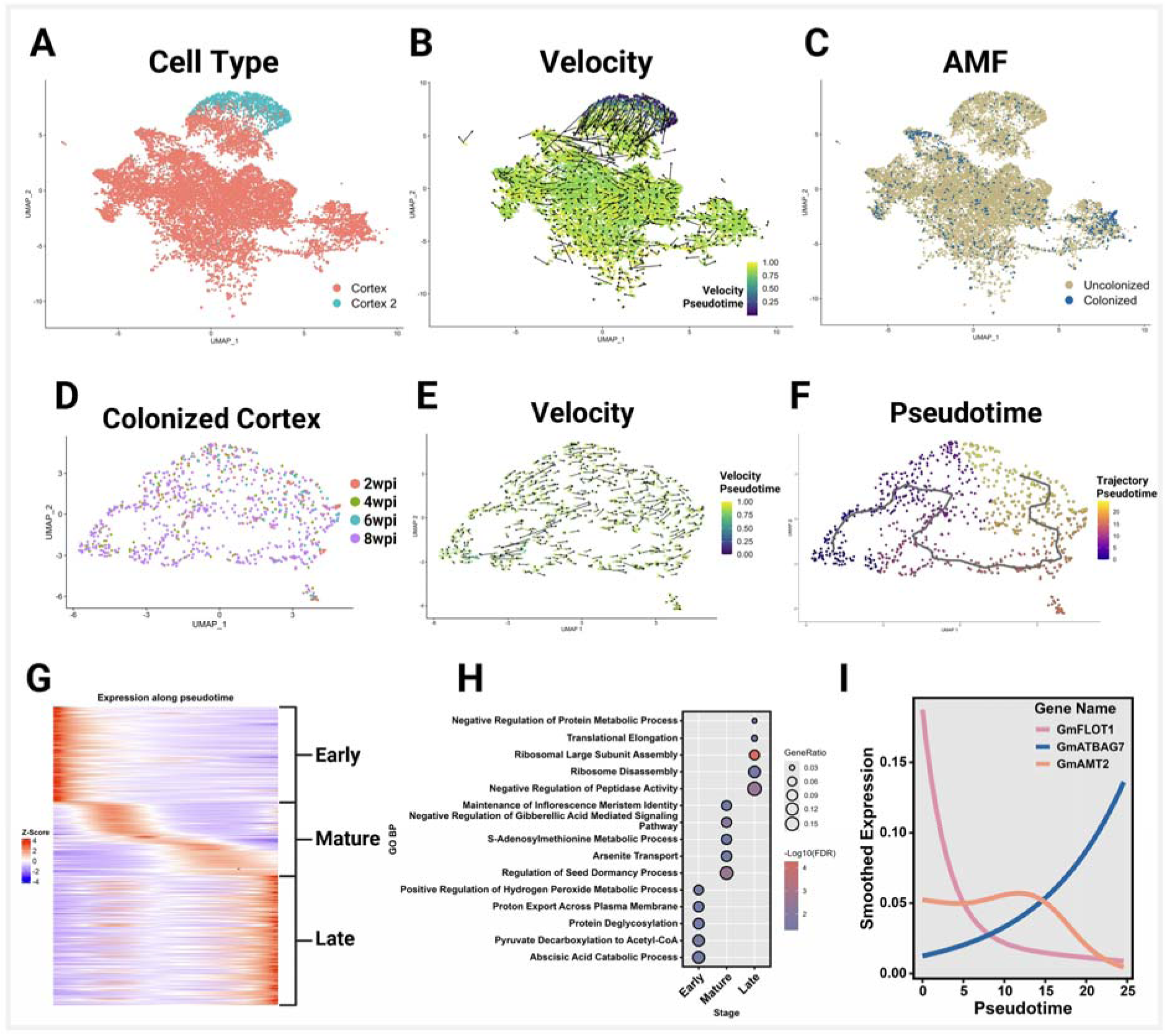
Cortex localized changes at the site of symbiosis. UMAP plots depicting; **(A)** cortex and cortex re-clustering (color represents cell type), **(B)** velocity pseudotime and trajectory (color represents velocity pseudotime and arrow direction indicates splicing trajectory), and **(C)** colonized vs uncolonized cortex cell distribution (color indicated colonization status). Colonized cortex nuclei UMAP plots depicting; **(D)** colonized cortex re-clustering (color represents sample), **(E)** colonized cortex velocity pseudotime and trajectory (color represents velocity pseudotime and arrow direction indicates splicing trajectory), and **(F)** colonized cortex monocle3 pseudotime and trajectory (color represents pseudotime and line indicates pseudotime trajectory). **(G)** Heatmap consisting of pseudotime DEGs plotted over pseudotime (x axis). Labelling on right to indicate early, mature, and late-stage expression programs. Color intensity represents scaled average expression (blue = low, red = high). Full gene list present in table S5. **(H)** Dotplot representing GO BP term enrichment of genes from groups in **(G)** (x axis). Top 5 GO terms based on fold enrichment plotted (p_adj_ < 0.05), color intensity represents significance (blue = low, red = high), dot size represents the ratio of detected genes to pathway genes in each GO term. **(I)** Line plot depicting representative gene from each group. X axis represents pseudotime and y axis represents smoothed expression from **(G)**. Color indicates genes as indicated in the legend.

Furthermore, to reconstruct cortical cell developmental trajectories during AMF symbiosis, we coupled pseudotime inference with RNA velocity, using unspliced-to-spliced mRNA transcript ratios as a directional readout of predicted future transcriptional state of each nuclei. The RNA-velocity analysis showed a conserved developmental path and was reinforced by pseudotime inference, which followed the same left to right cell fate trajectory (Fig. 5e, f). Leveraging this, negative binomial generalized additive model (NB-GAM) was fit to each gene using the tradeSeq framework [29], and genes showing significant pseudotime-dependent expression (FDR < 0.05) were grouped into early, mature, and late expression clusters and visualized on a heatmap (Fig. 5g, Table S5). GO term enrichment revealed temporally ordered programs across these stages in which early-stage arbusculated cells were enriched in *abscisic acid catabolism*, *hydrogen peroxide regulation*, and *proton export across the plasma membrane*. Mature-stage cells showed enrichment in *GA-mediated signaling*, whereas late-stage cells were enriched in *ribosome disassembly* and *negative regulation of protein metabolic process* (Fig. 5h). *GmFLOT1*, a lipid raft scaffold protein, peaked early consistent with periarbuscular membrane nanodomain establishment during initial colonization; *GmAMT2* peaked at the mature stage, reinforcing ammonium transport as the dominant N exchange activity at peak arbuscule function; and *GmATBAG7*, a BAG-domain autophagy co-chaperone, increased progressively and peaked late suggesting arbuscule dismantling and cellular recycling following symbiotic exchange (Fig. 5i).

### Combinatorial functional validation reveals a novel metabolic GO-switch for AMF symbiosis

Among the co-expression networks identified across the snRNA-seq atlas, the red network emerged as most spatially enriched in cortex 2 cells at 2wpi and 4wpi (Fig. 6a, S5, Table S3). GO term enrichment of red network genes revealed a significant enrichment in phenylpropanoid and coumarin biosynthesis and metabolism, indicating secondary metabolite reprogramming of cortex 2 cells during early colonization (Fig. 6b). Importantly, a hub gene, *GmF6’H1-2*, encoding a feruloyl-CoA 6’-hydroxylase that catalyzes a rate-limiting step in scopoletin-type coumarin biosynthesis showed high expression confined to 2wpi and 4wpi cortex 2 cells and expression in 2wpi endodermis (Fig. 6c) [36]. This expression pattern was shared among coumarin-related genes within the red network but was distinct from the broader expression patterns of other phenylpropanoid pathway genes (Fig. S15), indicating that the coumarin branch of the pathway is selectively activated in cortex 2 cells during early AMF colonization. Among all coumarins detected across the experiment, dimethylfraxetin was the only coumarin detected that exhibited differential enrichment in colonized segments at both time points (Fig. S16). Furthermore, both GmPAL2-2, a phenylalanine ammonia-lyase gene previously identified AMF colonization QTL on Chr. 20, and GmPAL1-4, located within a significant nodule number QTL on Chr. 09, (Fig. S15) showed cortex 2 specific expression patterns [37]. To directly test the functional role of *GmF6’H1-2* for AMF colonization, we leveraged whole-genome resequencing data to identify natural loss-of-function variants [37]. Six accessions (among ∼1,200 lines) carrying a shared two-base-pair deletion in the second exon of *GmF6’H1-2* were identified and confirmed by fragment analysis (Fig 6D, S17). This frameshift mutation introduces a premature stop codon at the 227th amino acid (S227*), leading to a truncated protein and thus constituting a loss-of-function allele (Fig. 6E).

**Fig. 6.**
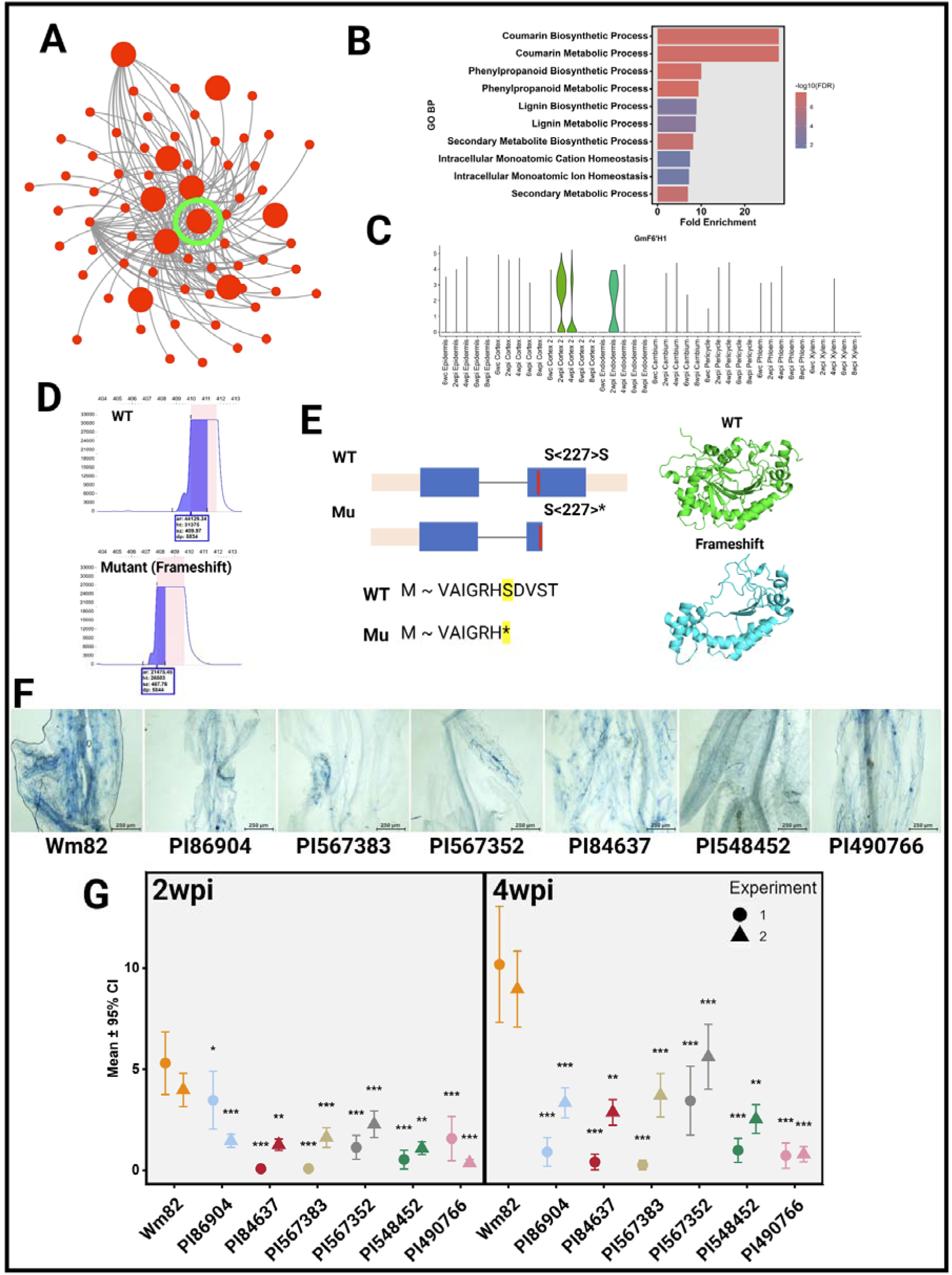
Coumarin pathway and hub gene validation. **(A)** visual representation of the red GCN specific to cortex cells. **(B)** GO BP term enrichment of red network genes. Full gene list of all networks presented in table S3. **(C)** Violin plot of *GmF6’H1-2* hub gene. Height of lines represents scaled expression level. Size of violin represents distribution of expression within the nuclei of the cell-type sample comparison groups; wider regions indicate a greater density of cells at that value. **(D)** Identification of mutations in *GmF6’H1-2* gene in selected lines using fragment analyzer and compared with Wm82. Full fragment analyzer results and other details for each soybean PI accession presented in Fig. S17. **(E)** Representative cartoon of frameshift mutation and location of amino acid change. Amino acid 227 changes from serine to stop codon, with the modelled WT and frameshift proteins present. **(F)** Representative trypan-blue stained root images of Wm82 and each frameshift soybean accession. Scale bars = 250 µm. Complete AMF colonization data is presented in table S6. **(G)** Mean ± 95% confidence intervals are shown for each genotype across two independent experiments. Colors denote genotype and shapes denote experiment replicate. Within each experiment and time (2wpi & 4wpi) group, Dunn’s post-hoc tests were used to compare each genotype against its corresponding Wm82 control. Significance levels are shown above points: * p < 0.05, ** p < 0.01, *** p < 0.001.

When all six natural variant lines and the reference genotype Wm82 were grown under identical AMF inoculation conditions, the six *GmF6’H1-2* loss-of-function accessions exhibited a consistent and significant reduction in AMF colonization percentage at both 2wpi and 4wpi relative to Wm82 (Fig 6F, G; Table S6). Collectively, these findings reveal a previously uncharacterized coumarin-centered metabolic GO-switch that is essential for the early establishment of AMF symbiosis in soybean. The convergence of co-expression network topology, hub gene specificity, spatial metabolomic validation, QTL co-localization, and natural loss-of-function genetics establishes *GmF6’H1-2* as a key regulator governing the transition from fungal recognition to successful cortical colonization.

## Discussion

### Integrated analysis enables cell-type and spatially resolved discovery in AMF symbiosis

AM symbiosis is among the most ancient and widespread plant-microbe interactions on Earth, yet the cell-type-specific regulatory logic governing each stage of colonization has remained poorly resolved, largely because bulk transcriptomic approaches conflate the transcriptional identities of the arbusculated cortex cell, the immune-activating epidermis, the MIR-priming pericycle, and every other root tissue into a single averaged signal [38]. By profiling 33,410 high-quality nuclei across four colonization timepoints and an uninoculated control, the snRNA-seq atlas generated here provides a breadth of genomic coverage for plant symbiosis studies, establishing a mechanistic framework of sufficient depth to resolve rare cell states, paralog-level gene expression differences, and dynamic colonization-stage transitions that would be invisible in previous datasets (Fig. 7). The identification of 20 distinct clusters spanning all principal root tissue layers is consistent with the known radial organization of the soybean root but extends it with single-cell resolution (Fig. S1a) [20–23]. We identified the cortex 2 cluster as a transcriptionally specialized population occupying a distal UMAP position to conventional cortical cells. Its disproportionate expression of cortex marker genes, combined with RNA velocity trajectories placed it upstream of the mature cortex, suggesting cortical progenitor population that progressively acquires symbiotic identity during colonization (Fig. 5A; S1B). As discussed below, this distinction matters mechanistically, because cortex 2 is the exclusive site of coumarin biosynthesis activation. Furthermore, the integration of spatial metabolomics across matched root sections adds a critical biochemical dimension to the transcriptional atlas. Because AMF colonization is spatially heterogeneous, not all segments within a root are colonized at any given timepoint. To support this, pixel-level metabolite profiling enabled colonized-versus-noncolonized comparisons within the same tissue section, providing an internal control that eliminates confounders such as plant age, P status, and developmental stage. The clear UMAP divergence of 2wpi and 4wpi colonized pixels from the uninoculated control, and from each other (Fig. 2B), demonstrated that AMF symbiosis imposes a progressive and stage-specific metabolic reprogramming rather than a static biochemical shift, and that this reprogramming is chemically distinct from P starvation responses in the control.

**Fig. 7.**
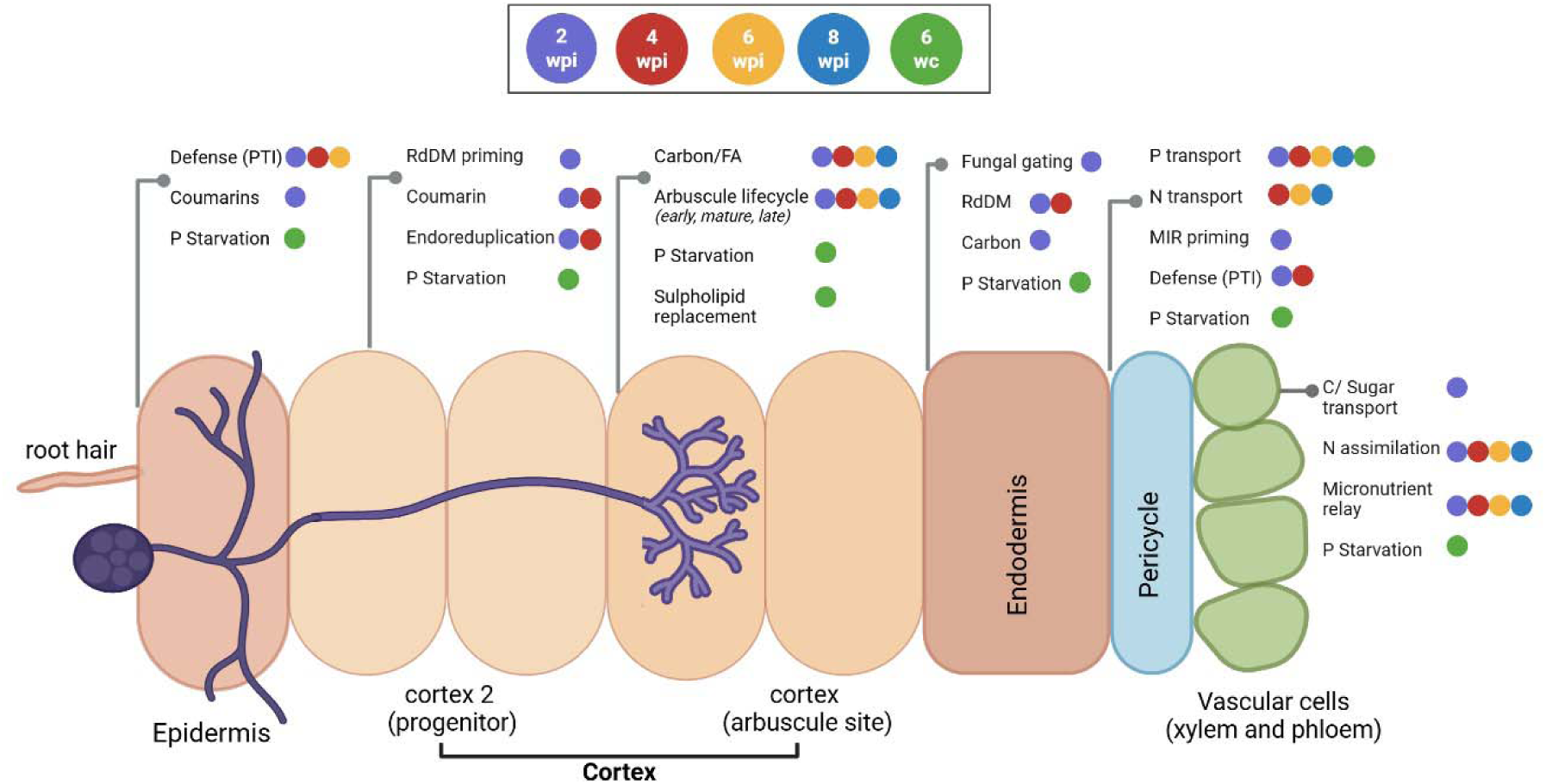
Proposed cell-type-resolved model of AMF symbiosis in soybean roots. The model integrates snRNA-seq and spatial metabolomics data to depict the spatially stratified, cell-type-specific molecular programs governing AMF colonization. Each root tissue layer contributes a distinct and non-redundant function: (i) the epidermis deploys a full PTI immune response that is temporally gated and progressively suppressed as colonization advances; (ii) the cortex 2 population activates a cortex 2-specific coumarin biosynthesis program (GmF6’H1-2 hub) and subsequently undergoes RdDM-mediated epigenetic silencing of NBS-LRR to permit arbuscule accommodation, accompanied by endoreduplication membrane remodeling; (iii) the mature cortex sustains arbuscule function through carbon and FA transport and transitioning to controlled arbuscule lifecycle and cell wall restoration; (iv) the endodermis provides a passive-active barrier for fungal gating and epigenetic priming; (v) the pericycle integrates AMF-delivered P and N into the vasculature through symbiosis-specific transporter repertoires; and (vi) the vascular cells mediates increased sucrose supply to colonized zones and systemic micronutrient relay. The P starvation was active throughout the root development in 6 weeks control roots.

The spatial metabolomics data revealed that AMF symbiosis in soybean proceeds through a biochemically coherent two-phase transition, spatially constrained to individual root segments, that directly mirrors the heterogenous biological progression of colonization (Fig. 2c). In early colonized tissue at 2wpi (cluster 4), the dominant metabolite classes were flavonoids and terpenoids. This metabolic state was mechanistically consistent with the established roles of flavonoids as pre-symbiotic host signals that stimulate AMF spore germination and hyphal branching, and of terpenoids as modulators of early immune responses and membrane remodeling during the accommodation phase of fungal penetration (Fig. 2d, e) [25, 39–42]. The co-enrichment of organic acids and amino acids at this stage suggested that broad priming accompanied the recognition and early accommodation of the fungal partner, potentially supplying energy for the membrane/cell wall remodeling that pre-penetration apparatus formation demands.

By 4wpi (cluster 8), the metabolic landscape shifts decisively in which free fatty acids became the enriched class, flavonoid enrichment declined, and phenolic acids and alkaloids increased (Fig. 2d, e). This transition was mechanistically significant because as documented before, AMF are obligate biotrophs that lack *de novo* fatty acid synthesis and are entirely dependent on host-exported lipids for membrane biogenesis and energy metabolism [40]. The emergence of a lipid-dominated metabolic environment at 4wpi, coinciding with established arbuscule function, is reflective of the core metabolic function of the mature symbiosis. Phenolic acid and alkaloid increase at this stage were consistent with sustained MIR priming rather than acute defense execution, as AMF colonization is known to elevate these compound classes systemically as part of a primed but not fully activated defense state [39]. The spatial resolution of this transition across individual root segments, and their absence from 4wc root segments, establishes that these metabolic shifts are driven by fungal presence rather than systemic developmental or nutritional cues.

### AMF induces active defense modulations at the cellular level

Plants face a fundamental immunological paradox when associating with AMF in which chitin and β-glucan in AMF cell walls trigger the same innate immune responses activated by pathogenic fungi, yet the plant must permit colonization while defending against true invaders [43, 44]. Compounding this, suppression of both SA and JA defense hormones is required within the colonization zone to permit expression of symbiosis-essential genes, yet JA-insensitive mutants show reduced symbiosis and AMF has been shown to induce SA biosynthesis in the host [45–47]. Resolving these simultaneous opposing programs requires cell-type resolution, as bulk transcriptomics inevitably merge the immune-suppressing cortex with the immune-activating epidermis. The snRNA-seq atlas resolves this paradox through a three-zone spatially stratified immune architecture. At the epidermis, 2wpi cells mounted the canonical and complete PTI response, encoded entirely within the brown co-expression network. This network span receptor complex activation (*GmLIK1, GmBIR1*) [48, 49], the MAPK cascade (*GmMEKK1, GmMPK3, GmMKP1*) [50], and RBOH-driven ROS burst (*GmRBOHB, GmRBOHD*) [51], with parallel SA amplification via the EDS1-PAD4-SAG101 complex [52] and JA initiation through *GmLOX/AOS/AOC* [53] (Fig. 3a, S6, Table S3). Critically, *GmNSL1*, a MACPF-domain negative regulator, potentially preventing this cascade from escalating into hypersensitive cell death [54], positioning the epidermis as a temporally gated immune checkpoint rather than a committed defense executor (Table S3). This epidermal PTI response declined progressively to control levels by 8wpi, suggesting that AMF-induced suppression of defense is integrated as colonization becomes established. In the pericycle, a distinct MIR program operated in parallel. In 6wc (control) roots, *GmJAR1-2/3* and *GmCOI1-1/4* route JA-Ile toward growth regulation rather than MIR priming [55, 56]. By contrast, 2wpi pericycle cells express *GmJAR1-1/4, GmMYC2-1/2/3*, and *GmERF2-2* without COI1 or MKK3 repressors, enabling MYC2 accumulation and vascular MIR priming [55, 56]. At the endodermis, miRNA processing enrichment alongside casparian strip reinforcement constitutes a dual passive-active barrier preventing fungal overgrowth into the vasculature (Fig. 3a) [11].

Spatial metabolomics spatially supported this transcriptional architecture. Defense-related coumarins accumulated in the outer (epidermal) colonized zone at 2wpi [57], while proanthocyanidins and others concentrated in the inner colonized zone, mirroring the brown network’s MVA pathway flux (JAZ1 degradation to *GmHMG1* to triterpene/saponin synthesis [58, 59]) and *GmMYB14/15*-driven proanthocyanidin biosynthesis [60]. A second metabolite group consisting of plumeranes, quinorisidine, phenolic acids, and diterpenoids maintained elevated levels in both 2wpi and 4wpi colonized segments. This is mechanistically consistent with MIR priming sustained by *GmOPCL1/GmACX1* modulating steady JA flux and *GmERF1/3/5/9* providing the JA/ethylene co-signaling arm [61–63]. Biflavones and terpenoid alkaloids appeared exclusively at 4wpi, reflecting a likely precursor accumulation requirement [64, 65]. Critically, none of these defense or MIR metabolites were detected in noncolonized tissues, establishing AMF colonization and not P starvation as the necessary inducing condition (Fig. 3a). Finally, the near-complete absence of defense gene expression in colonized cortex cells indicates deliberate epigenetic immune suppression rather than passive tolerance. The RdDM pathway genes *GmNRPD2A-1/2, GmAGO4-2/4/5, GmET2*, and *GmCMT3-1/2* were specifically induced in colonized cortex cells at 2wpi [66], encoding the Pol IV/Pol V siRNA biogenesis and AGO4-facilitated silencing machinery that targets NBS-LRR defense loci [67] (Fig. 3a, S3, S4, S14). In soybean, RdDM downregulation during pathogen infection relieves NBS-LRR silencing to permit defense gene transcription, but the inverse pattern is observed here, wherein RdDM upregulation with concurrent NBS-LRR suppression in AMF-colonized cells, positions this pathway as the cortex-specific epigenetic accommodation mechanism [68] (Fig. S3, S4). Co-induction of *GmCMT3-1/2* further established a self-reinforcing methylation loop that propagates silencing beyond the initial transcriptional window, explaining how the 2wpi expression spike is sufficient to maintain defense suppression across all subsequent colonization stages (Fig. S3c) [66]. Upstream of this, broader induction of *GmHSC70-1/2/5* and *GmHSP90.4* paralogs across both colonized and uncolonized cortex cells indicates AMF-induced pre-loading of the AGO4 chaperone complex prior to local RdDM activation, suggesting a priming mechanism that anticipates fungal contact at the tissue level given the relatively slow kinetics of RdDM (Fig. S14b) [34, 69].

### AMF symbiosis shapes plant carbon dynamics

AMF colonization restructures plant carbon allocation by establishing the root cortex as a dominant metabolic sink consuming approximately 20% of plant-fixed photosynthates [4]. The snRNA-seq data reveals a two-zone sucrose catabolism that utilizes carbon from multiple routes simultaneously. Elevated *GmSUC2-2/3/6* expression in root phloem at early colonization timepoints reflects increased phloem transport capacity to meet fungal carbon demand, with *GmSUC2-6* absent from the 6wc control indicating AMF-specific recruitment of additional transport capacity (Fig. S7a) [70]. Following phloem unloading, *GmATSUS4-1/2* are induced mainly in 2wpi cortex 2 cells, the primary symbiosis interface region, while *GmATSUS4-3/4* are simultaneously induced in xylem, enabling radial sucrose cleavage from the vascular supply route (Fig. S7b) [71]. The AMF-specific induction of both paralog pairs at 2wpi, coinciding with the onset of arbuscule formation and peak fungal carbon demand, is consistent with a direct transcriptional response to the fungal sink rather than constitutive sucrose catabolism [72, 73]. Co-induction of *GmCINV2-1/2* and *GmA/N-InvC-2* in 2wpi xylem cells likely reflects sucrose catabolism channeled toward cell wall reinforcement and root growth to accommodate the expanding fungal network (Fig. S7b) [74]. The glucose produced by these pathways feeds the periarbuscular membrane hexose efflux required for arbuscule maintenance, as demonstrated by the requirement of SWEET1b-class transporters for arbuscule integrity in *Medicago* [75]. Spatial metabolomics validates this two-zone model wherein maltotriose from amylase-driven starch hydrolysis accumulates in colonized middle and outer zones (cortex route), while acetyl-maltose and D-maltotetraose accumulate in colonized inner and middle zones (vascular route), with the spatial complementarity of these glucose polymers directly mirroring the two anatomical catabolism zones identified in snRNA-seq (Fig. S8b) [76]. This observation is consistent with xylem-localized sucrose breakdown generating poly-glucose intermediates through the *GmATSUS4-3/4* route (Fig. S7, S8b) [77, 78].

The fatty acid (FA) biosynthesis reprogramming in colonized cortex cells constitutes the mechanistic core of AMF carbon provisioning [79, 80]. Rather than broad upregulation of lipid biosynthesis, AMF fundamentally reprograms plastid identity in cortex cells, to maximize C16:0 production [81]. The co-induction of the complete FASII module, from malonyl-CoA committed synthesis (*GmCAC1, GmMTKAS*) through acyl carrier protein cycling (*GmACP3/4, GmmtACP2/3*) to condensation enzymes (*GmKASI-1/2*) and enoyl reduction (*GmMTHD-1/2*), in 2wpi cortex cells represents the transcriptional signature of arbusculated cell differentiation to supply the RAM2/STR2-mediated export pathway to AMF (Fig. 4a, S8a) [30, 82]. The temporal shift to *GmFATB* paralog expression at 4–8wpi marks the transition from de novo FA assembly to actively exporting it as colonization matures (Fig. S8a) [83]. This staged reprogramming of FASII assembly at 2wpi followed by FatB-driven export maintenance, mirrors the fungal carbon dependency established in *Medicago*, where FatM thioesterase and RAM2-facilitated β-monoacylglycerol production are required specifically in colonized cortex cells [30, 84].

Spatial metabolomics further confirmed this temporal changes as C13–C20 long-chain lipids were elevated in 2wpi colonized segments with a reduced but sustained signal at 4wpi, while methyl stearate and methyl ricinoleate accumulated specifically in colonized middle cortex at both timepoints, providing individual lipid species-level validation of active plastidial FatB flux (Fig. 4b, c, S8b) [30]. In the late stage cortex cells *GmKCS* paralogs likely represent metabolic overflow rather than a primary export program wherein KCS elongases extend C18:0-CoA when FASII flux exceeds FATB export and phospholipid assembly capacity (Fig. S8a) [85, 86]. Very-long-chain lipids (C21–C34) accumulated specifically in noncolonized rather than colonized segments, highlighting that in colonized cells the C16:0 pool is continuously utilized by RAM2/STR2 for transfer to the AMF (Fig. 4b, c) [86]. Membrane lipids LysoPC and oxidized phosphatidylcholine species in colonized segments further evidenced active periarbuscular membrane biogenesis at the plant-fungus interface (Fig. S8b) [87, 88].

### Cellularly resolved nutrient dynamics in response to AMF symbiosis

Nutrient uptake and transport are key outcomes of plant–microbe symbiosis and defining their molecular and cellular mechanisms is essential for crop improvement. This study revealed a temporally progressive repression of the canonical P starvation response as AMF colonization matures. In 6wc plants, key P starvation genes *GmPHR25* [31] and *GmPHR1-3* were broadly expressed across epidermal, endodermal, cambium, and pericycle cell types, with *GmSIZ1-1* (the SUMO E3 ligase amplifying PHR1 output) peaking in 6wc endodermis (Fig. S12a) [89]. Unlike most species where PHR1 is constitutively expressed, soybean PHR homologs are P-status responsive [32, 90], and *GmPHR1-1/2/3* declined progressively from 2-4wpi to undetectable levels by 6-8wpi, directly tracking AMF-mediated P sufficiency restoration. Notably, *GmSPX2-5* and *GmSPX3-1/2*, the inositol pyrophosphate Pi sensors that suppress PHR1 under P sufficiency, appeared transiently in 2-4wpi cortex 2 cells then suppressed by 6wpi, while in 6wc plants the same SPX genes reached maximum expression without achieving PHR1 suppression. This is consistent with SPX domain loading being blocked under chronic starvation [88], establishing that AMF colonization restores the Pi-sensing competence that P starvation itself impairs. We observed a clear functional outcome of this circuit, wherein, the downstream P-scavenging genes NPC4 (hydrolyzes membrane phospholipids to release P) and *GmSQD2* (substitutes phospholipids for sulfolipids to reduce cellular P demand) were highly expressed in 6wc cell types and were entirely absent from all AMF-colonized timepoints (Fig. S12a) [32, 90]. Their complete withdrawal from colonized tissue confirms that AMF-delivered P is sufficient to suppress the PHR1 program by mid-colonization, and that the metabolic cost of membrane lipid remodeling under P stress is averted by successful symbiosis.

The PHO1/PHO2 vascular P loading families exhibited a paralog-level functional segregation between starvation-driven and symbiosis-specific P export not previously described in soybean. Under canonical P starvation, PHR1-induced miR399 degrades PHO2 mRNA, relieving PHO2-mediated ubiquitination of PHO1 and enabling xylem P loading [91]. Here, *GmPHO1-2/3* and *GmPHO2-2/3* operated in both 6wc and AMF-treated pericycle, consistent with shared starvation and symbiosis roles. However, *GmPHO1-1/4* and *GmPHO2-4* showed exclusive AMF-induced pericycle expression from 2wpi onward, entirely absent from 6wc tissue, indicating that symbiosis recruits a distinct transporter repertoire beyond what starvation alone activates (Fig. S12b) [92]. This functional divergence has precedent in PHO1 specialization in *Vigna radiata* and PHO2 in *Medicago truncatula* [93–95] and aligns with the AMF-specific carbon transport gene-duplicate recruitment described above, suggesting paralog-level functional specialization is part of soybean symbiotic nutrient loading, however this inference needs functional validation.

On the other hand, N mobilization operated through two coordinated zones. *GmGLT1-1* induction in 2-4wpi cortex 2 and *GmGLT1-2/3/4* induction in AMF-treated xylem cells established a biphasic GOGAT ammonium assimilation program absent from 6wc tissue, consistent with documented GOGAT upregulation in response to AMF-delivered ammonium (Fig. S12c) [96]. Induction of *GmCEPR1-1/2* in AMF-treated phloem cells is most likely responsive to elevated available saccharide levels, and the concurrent AMF-specific induction of sucrose transport, sucrose catabolism, and poly-glucose accumulation detected in this dataset provides an induction stimulus independent of N status (Fig. S8b, S11) [97]. The downstream NRT expression, *GmNRT2.4-1/3/4* and *GmNRT3.1-1/2*, in epidermal cells concurrent with elevated CEPR1 is consistent with the known link between NRT gene activation and CEPR1 activity (Fig. S12b) [98]. The coordinated upregulation of P and N transport programs identified here provides a transcriptional basis for the preferential aboveground biomass allocation observed under combined symbiosis. In our previous study, combined AMF and rhizobia treatment produced a near-double aboveground biomass without a corresponding increase in root mass [37]. This would suggest that once symbiotic nutrient delivery is established, the plant redirects assimilated N and P toward shoot growth rather than continued root expansion, a resource allocation shift now traceable to the pericycle-centered nutrient loading programs identified in this dataset. Beyond P and N, AMF-induced expression of copper, molybdate, chloride, manganese, and boron transporters (primarily in pericycle and phloem cells) established that symbiotic nutrient enhancement extends to broad micronutrient acquisition, with the pericycle functioning as the central integration hub and phloem as the systemic relay to distal tissues (Fig. 4d) [37, 99].

### Cortex localized changes at the site of symbiosis

The cortex is the primary site of arbuscule formation [100], but its transcriptional heterogeneity (colonized vs non-colonized cells) is obscured in whole-root analyses. Focused sub-clustering of cortex and cortex 2 cells reveals that these populations are connected across a developmental stage, with RNA velocity trajectories establishing cortex 2 as a transcriptionally upstream, less differentiated population that progressively gives rise to the cortex cell states engaged in symbiosis (Fig. 5). This developmental relationship between cortex cells has not previously been described and provides a cellular lineage framework for interpreting the stage-specific gene programs identified throughout this study. For example, the coumarin biosynthesis sub-program is a cortex 2-localized early event, while the FASII lipid reprogramming, arbuscule-maintenance, and RdDM programs represent later cortex developmental state activities.

Marker-guided identification of colonized vs uncolonized cortex cells revealed two processes exclusive to colonized cells (Fig. 5c). First, cell cycle, DNA replication, endoreduplication, and DNA repair GO terms are strongly enriched in 2wpi colonized cortex cells (Fig. S14c). This restriction to early colonized cortex is consistent with AMF-induced endoreduplication in *Medicago* as a CSSP-dependent pre-penetration response spatially confined to arbusculated and immediately neighboring cells [35]. This pattern indicates that endoreduplication by amplifying the genome without cell division provides the transcriptional capacity to sustain the cell growth, differentiation, and metabolic specialization necessary for arbuscule accommodation. Vesicle trafficking revealed a complementary architecture (Fig. S14d). Endocytosis related genes dominated at 2-4wpi colonized cortex, aligned with clathrin-mediated endocytosis in *Medicago* colonized cortex through AP2A1/CHC2/TPLATE co-induction, and with endocytic uptake of fungal-derived extracellular vesicles in the periarbuscular space [101, 102]. By 6-8wpi, the overriding signal shifted to exocytosis, vesicle fusion, and vesicle-mediated transport in both colonized and uncolonized cortex cells, consistent with VAMP72-class v-SNAREs and exocyst-promoted PAM delivery sustaining the mature interface during peak nutrient exchange (Fig. S14d) [103].

The pseudotime trajectory of colonized cortex cells reconstructed the complete arbuscule lifecycle, from initial accommodation through nutrient exchange to controlled senescence within a single continuous cell population irrespective of the plant age, capturing biological transitions of the arbuscule lifecycle that typically span 3 to 8 days (Fig. 5d, e, f) [104]. This resolution has not previously been reported across a multi-timepoint series in a legume host. Based on GO term analysis, at early pseudotime, ABA catabolism (via *GmCYP707A3*) reflects the documented requirement to suppress ABA locally to permit colonization to proceed [105, 106], while pyruvate decarboxylation to acetyl-CoA channels carbon (via *GmPDH-E1 ALPHA*) suggest FASII burst to supply the fungus with lipid at the onset of colonization (Fig. 5g, h, Table S5) [107]. Transient *GmFLOT1* expression organizes periarbuscular membrane lipid raft microdomains for symbiosis [108], while concurrent *GmATB1* induction provides a second, RdDM-independent layer of cell death suppression (Fig. 5i, Table S5) [109]. Together, these mechanisms ensure membrane assembly and immune gating operate simultaneously at the early timepoint of fungal contact. The mature stage, marked by *GmAMT2* expression, captures the functional nutrient exchange window and enrichment of *S-Adenosylmethionine Metabolic Process* and not previously reported in AMF datasets (Fig. 5h, i). SAM cycle activity is mechanistically required to sustain the methyl donor flux powering ongoing RdDM-mediated NLR silencing in the same cells, suggesting that immune suppression maintenance is an active metabolic cost of the mature arbuscule state (Table S5) [110]. Concurrent enrichment of *Negative Regulation of Gibberellic Acid Mediated Signaling Pathway* reflects the established GA-symbiosis antagonism in which gibberellin suppression is required to sustain arbuscule integrity (Fig. 5h) [111].

At late pseudotime, the transcriptional program shifts from nutrient exchange to cellular dismantlement. The near-complete collapse of *GmAMT2* and *GmFLOT1* expression suggests functional inactivation of the arbuscule wherein ammonium import and the periarbuscular membrane nanodomains that scaffolded nutrient transfer are disassembled (Fig. 5i). Concurrent induction of the ER-localized co-chaperone *GmATBAG7* [112], suggests that the cell is actively managing the PAM system dismantling rather than passively decaying. Similarly, ribosome activity at this stage is highly essential and we observed co-enrichment of *Ribosomal Large Subunit Assembly* alongside *Ribosome Disassembly* and *Negative Regulation of Peptidase Activity* that indicates a tightly bound burst of selective protein synthesis before global translational shutdown (Fig. 5h) [113]. Kunitz Trypsin Inhibitor paralogs (*GmKTI5*) appeared to pace this window by suppressing premature proteolytic degradation of the translation apparatus, ensuring that late-stage gene products are still produced before ribosome turnover proceeds (Table S5) [114]. Simultaneous induction of primary cell wall related genes *GmCESA6*, *GmXTH9*, *GmEXT3*, and *GmPRX52* constituted a sequential wall loosening, resynthesis, and stiffening program that restores cortex cell wall integrity following arbuscule collapse [115–117], completing the arbuscule lifecycle and returning the cell to a wall-intact, functionally-competent resting state (Table S5).

The late-stage green co-expression network, active exclusively in 6-8wpi cortex cells and absent from 6wc, encoded the more widespread arbuscule molecular program. First, ROS homeostasis at the mature arbuscule interface is managed through a copper-centered dismutation axis, in which *GmATX1* delivers Cu to the superoxide dismutase *GmCSD1*, while metallothionein paralogs buffer excess Zn²⁺/Cu²⁺ to prevent metal-catalyzed oxidative damage (Fig. S13, Table S5) [118–121]. PIP aquaporins likely facilitate H O partitioning across the periarbuscular membrane, though their precise role in redox compartmentalization at this interface warrants further investigation [123–126]. Second, co-expression of GTPases (*GmARFA1F/GmRAB6A*), SNARE complexes (*GmVAMP726/GmSYP51*), together with lipid remodeling genes (*GmPLD*α*1/GmACBP6/GmDGAT3)*, sustains continuous membrane biogenesis to support arbuscule branching and expansion as the symbiotic interface [33, 122–124]. Third, ubiquitin-SCF complex components and senescence-associated proteases provided controlled protein turnover as arbuscule branches begin to senesce (Table S5) [125]. The complete absence of the green module from uninoculated 6wc roots confirms that this is a colonization-specific maintenance program, not a generic stress response, and distinguishes it mechanistically from the early epigenetic gating and mid-stage nutrient exchange programs described above (Fig. S7).

### Coumarin is functionally imperative for effective AMF colonization and establishment

The co-expression network analysis identified the red network which was exclusively active in cortex 2 cells at 2-4wpi (Fig. 6a, S7). This network encoded a biochemically complete coumarin biosynthesis-to-secretion pathway whose hub architecture, spatial metabolite accumulation, GWAS convergence, and natural loss-of-function phenotype collectively constitute four independent lines of evidence for a previously uncharacterized pro-symbiotic function in a plants (Fig. 6, S15, S16, S17) [37]. Within the network, hub status is limited to the committed entry enzymes (*GmF6*′*H1-1/2*), the rhizosphere export transporters (*GmABCG37-1/2*), and the scopoletin O-glucosyltransferase (*GmCOSY*) (Table S3). The intervening biosynthetic steps *GmC4H-2, GmC3H-2, GmCCoAOMT1-2, Gm4CL2-2* supplying feruloyl-CoA, and *GmS8H-1, GmOMT1* completing the scopoletin-to-dimethylfraxetin conversion, occupy non-hub network positions (Fig. S15, table S3) [28, 36, 126, 127]. This network architecture is informative because the transcriptional co-regulation was most tightly organized around the flux control nodes of the pathway (the committed entry enzyme and the export transporter), while the intermediate steps were co-expressed but not co-regulated at the same level. The broader co-expression but non-network clustering of trunk *GmPAL, GmC3H, GmCCoAOMT*, and *Gm4CL* paralogs at 2-4wpi cortex 2 confirms that the phenylpropanoid supply chain upstream of the network boundary is co-activated but not incorporated into the specialized coumarin (red) module (Fig. S15). This line of evidence indicates a metabolic channel from the general phenylpropanoid pathway into the dedicated coumarin sub-pathway. Among all coumarins detected by spatial metabolomics, dimethylfraxetin was the sole compound showing colonized-segment-specific enrichment at both 2wpi and 4wpi. This identified dimethylfraxetin as the AMF-specific pathway endpoint and provides spatially explicit metabolite-level validation of the red network’s predicted output. This represents a cross-omics corroboration that connects the co-expression module to its chemical product in the tissue where that module is active (Fig. 6, S15, S16). The genetic significance of this pathway was independently reinforced by two significant GWAS candidate genes that mapped to the phenylpropanoid entry step wherein *GmPAL2-2* and *GmPAL1-4* associated with symbiosis were both clustered with red network genes in cortex 2 (Fig. S15). These results established that natural allelic variation at the pathway’s entry point is a genetically validated determinant of AMF colonization rate [37].

The functional validation of this pathway for colonization efficiency was tested by allele mining of the diverse germplasm collection for natural loss-of-function (Fig. 6d, S17). Six independent soybean accessions carried a confirmed 2-bp frameshift in *GmF6*′*H1-2* that introduced a premature stop codon at S227* and produced a truncated, non-functional protein. All mutant lines exhibited statistically significant reductions in AMF colonization percentage at both 2wpi and 4wpi relative to the reference genotype Wm82 (Fig. 6f, g, S27, Table S6). The phenotypic consistency across six independent genetic backgrounds carrying the same confirmed allele and two independent experiments, constitutes robust forms of functional validation available in a polyploid crop species. The mechanistic basis for collapsed colonization is likely multifactorial. Root-secreted scopoletin has been shown to stimulate *Rhizophagus irregularis* metabolism [128], suggesting that coumarin secretion from cortex 2 cells via *GmABCG37* may facilitate hyphal chemotropism or pre-penetration apparatus formation at the cortical colonization front (Table S3). Coumarin pathway overexpression can partially rescue AM incompatibility in non-host *Arabidopsis* [128], indicating that the pathway’s pro-symbiotic activity is transferable across species. Additionally, coumarins modulate local iron availability through chelation, and iron homeostasis has been implicated in periarbuscular space, suggesting that *GmF6*′*H1-2* loss-of-function may also impair the iron microenvironment required for efficient arbuscule establishment [127, 129].

The convergence of co-expression network hub architecture, spatial metabolomics co-localization of the pathway endpoint, GWAS corroboration, and natural loss-of-function genetics constitutes the comprehensive validation of a single symbiosis gene reported in soybean. The identification of *GmF6*′*H1-2* as a pro-symbiotic hub gene with validated natural loss-of-function alleles also opens a direct breeding route by screening germplasm panels. Identification of increased function *GmF6*′*H1-2* alleles and selecting for high-colonization accessions could enhance AMF-mediated nutrient acquisition, with potential applications for sustainable reduction of synthetic fertilizer inputs.

## Conclusion

This study provides the comprehensive cell-type and spatially resolved molecular atlas of AMF symbiosis in plants. By integrating snRNA-seq with spatial metabolomics on matched root sections, we resolve the cell-autonomous molecular logic of every major root tissue during fungal colonization (Fig. 7). The central organizing principle had emerged that AMF symbiosis is spatially stratified, temporally ordered, and cell-type-specific series of coordinated programs (Fig. 7). Immune gating at the epidermis (brown co-expression network), epigenetic accommodation in colonized cortex cells, a two-zone sucrose-to-lipid carbon provisioning, pericycle-centered nutrient loading, and a coumarin biosynthesis switch in cortex 2 cells collectively constitute the molecular architecture of effective AMF symbiosis in soybean. The pseudotime reconstruction of the arbuscule lifecycle from endoreduplication, periarbuscular membrane biogenesis, and peak ammonium exchange provided a continuous single-cell resolution description of a biological process previously inaccessible. Together, these findings reframed AMF symbiosis as an orchestrated multicellular program where each root cell type contributes to the establishment and maintenance of the symbiotic interface.

Beyond their mechanistic significance, these findings carry direct translational relevance. The identification of *GmF6’H1-2* as a validated pro-symbiotic hub gene with confirmed natural loss-of-function alleles that reduce colonization efficiency provides actionable breeding and biotechnological targets to enhance AMF colonization efficiency. Similarly, the cell-type-resolved P and N transport programs defined here, particularly the AMF-specific PHO1/PHO2 and NRT/AMT paralog pairs induced exclusively under symbiosis, provided putative candidate genes targeting reduced synthetic fertilizer dependency. More broadly, the multi-omic atlas described here serves as a high-resolution reference framework that can anchor comparative studies across legume species and inform synthetic biology efforts to transfer symbiosis competence to non-host crops. The atlas provides the transcriptional and metabolic baseline against which the effects of agronomic management, climate variation, and fungal strain diversity on symbiotic efficiency can be precisely measured. As soil health and sustainable nitrogen and phosphorus cycling become increasingly critical to global food security, a mechanistic understanding of the molecular architecture of this ancient plant-fungus mutualism at the cellular resolution is not merely academically valuable but practically essential.

## Materials and methods

### Plant material, microbial inoculation, and growth conditions

The soybean cultivar Williams 82 was used in this study. Plants were grown individually in 1-gallon pots, filled with commercial-grade coarse sand (Quikrete). Sand was sterilized by double consecutive autoclave cycles at 121°C. The composite AMF inoculant product used for this experiment was obtained from The International Collection of (Vesicular) Arbuscular Mycorrhizal Fungi (INVAM), The University of Kansas, USA. The inoculum was comprised of five arbuscular mycorrhiza strains (*Rhizophagus irregularis*, *R. clarus*, *R. intraradices*, *Funneliformis mosseae*, and *Claroideoglomus etunicatum)*. Approximately twenty grams of the AMF inoculum, containing 600-700 spores 100g^−1^, were applied to each pot of soybean. Soybean seeds were surface sterilized in 10% bleach and sown on top of the sand/AMF inoculum. Plants were grown under controlled greenhouse conditions and sample collection took place 2, 4, 6, and 8 weeks after emergence. For each timepoint, root tissue was pooled from 5 individual plants for snRNA-seq and for spatial metabolomics prior to nuclei isolation/cryosectioning. Pooling was necessary to obtain sufficient nuclei yield per 10X lane. To assess whether pooled samples adequately represent biological variation, per-plant colonization percentage (Trypan blue staining) was quantified prior to pooling.

### Root phenotyping

Plants were gently uprooted at the respective time points. Fresh root segments ∼1 cm in length were randomly collected from each plant for AMF colonization confirmation. The rest of the root was flash-frozen in liquid nitrogen for nuclei isolation and library preparation. The root segments were stained as outlined in D’Agostino, Raturi [37].

### Nuclei isolation, snRNA library preparation, and sequencing

Nuclei were isolated using an adapted protocol from Guillotin, Rahni [130]. Briefly, tissue was ground in a mortar and pestle in liquid nitrogen. Samples were transferred to a 1.5mL microcentrifuge tube with nuclei isolation buffer, and ground with a plastic pestle. Samples were then filtered on 100um, 70um, and 30um cell filters (Miltenyi Biotec). Samples were centrifuged and the nuclei pellet was resuspended in nuclei was buffer. The samples were centrifuged again and resuspended in nuclei final buffer. The nuclei were stained with propidium iodide and counted with the Luna-FL cell counter (Logos Biosystems). The nuclei were then loaded onto the chip as per the manufacturer’s recommendations to target 10,000 nuclei recovered (10X Genomics, Pleasanton, CA, USA). Library construction for Illumina sequencing was performed with the Chromium™ Single Cell 3’ Library & Gel Bead Kit v3.1 protocol (10X Genomics). The single-indexed sequencing of paired-end libraries were carried out on an Illumina™ NovaSeq 6000 platform according to the 10X Genomics manual.

### snRNA-seq data pre-processing, clustering, and cell type annotations

Sequencing data was processed using kallisto bustools [131] to obtain cell feature counts for analysis. The data was aligned to the Glycine max v4 genome. Count data was further analyzed within the R package Seurat v4 [19]. Chloroplast and mitochondrial reads were removed, and nuclei in the 95th percentile for gene count and 90^th^-99^th^ percentile for UMI were selected for further pre-processing. Then, doublets were removed with scDblFinder [132], and background RNA was removed with SoupX [133]. The data was then normalized by SCTransform, and the samples were integrated using Harmony [134] using PCs 1 to 30 to reduce batch effect and identify conserved cell types. Dimension reduction of the integrated data was performed using harmony reduction, and the UMAP plots were generated using dims 1 to 30. Monocle3 clustering was applied to cluster cells and identify the potential cell types [135]. Annotation of cell types was performed by utilizing the published soybean single-cell markers from (Fig. 1c, S2, Table S2) [20–23]. Expression of the markers was analyzed using the AddModuleScore function from Seurat and observed using the dotplot and featureplot functions.

### Spatial metabolomics processing and data analysis

Frozen root samples from 2wpi, 4wpi, and 4 week uninoculated control were processed for analysis. Briefly tissue was cryosectioned in slices of 18μm thickness. Slices were transferred to a pre-cooled indium tin oxide (ITO) coated glass slides and vacuum dried for 30 minutes. A 15 mg/mL solution of DHB (2,5-dihydroxybenzoic acid) was prepared in a solvent mixture consisting of 90%:10% Acetonitrile:water. This DHB matrix solution was evenly applied onto indium tin oxide (ITO) slides bearing tissue slices using a TM-Sprayer matrix sprayer, employing the following instrumental parameters: a temperature of 60°C, a flow rate of 0.12 mL/min, a pressure of 6 psi, a total of 25 gas cycles, and a drying time of 5 seconds between each cycle. Tissue section slides were subject to laser energy (50-1300 Da) and desorbed molecules were identified by the mass spectrometer as a resolution of 50μm^2^. Raw data were imported into SCiLS Lab software, with root mean square (RMS) normalization to obtain relative intensity information. Compound identification was conducted by aligning the collected spectra with entries in both the Metware Biotechnology database and public repositories using their primary molecular weights within a margin of error of 10 ppm. Per pixel (50μm^2^) metabolite abundance data was used to generate UMAP plots using the Seurat package. Clustering was performed similarly to the snRNA-seq data, via monocle3 [135]. Root segment visualization was plotted using ggplot2’s geom_tile function [136]. Radar plots were generated via the fmsb package [137]. Spatial regions were determined using the EBImage R package [138].

### snRNA-seq data analysis

Gene co-expression networks (GCN) were generated using the top 5000 genes expressed by the CSCORE package [139]. GCN representation was performed in cytoscape [140]. GO biological process term enrichment was performed using the topGO package in R using gene ontology terms obtained from SoyBase database [141]. Pseudotime trajectory analysis was performed using monocle3 and pseudotime DEGs were filtered using the tradeseq packages in R [135, 142]. Velocity trajectory analysis was performed using the scvelo package in R [143]. For cortex analysis, the cell types cortex and cortex 2 were subset from the whole Seurat object and reclustered using runUMAP [19]. The colonized cortex was identified using the AddModuleScore function with marker gene orthologs from Serrano, Bezrutczyk [13] (Table S4). Nuclei with the top 5% expression of the AMF marker gene module were selected as putative colonized cells, and subset again for further analysis. Expression of genes and pathways were visualized using the AddModuleScore function from Seurat and observed using the dotplot and vlnplot functions.

### F6’H1 natural variant experiment

Soybean accessions PI86904, PI84637, PI567383, PI567352, PI548452, PI490766 were chosen based on variant calling data generated in D’Agostino, Raturi [37] containing a 2bp deletion in the F6’H1, which was confirmed by fragment analyzer. Briefly, leaf tissue was collected from each genotype, and DNA was extracted using the CTAB method [144]. Gene specific primers were designed flanking the potential mutation site (Table S8). DNA fragments were analyzed via capillary electrophoresis on the SeqStudio Genetic Analyzer (Thermo Fisher Scientific). Electropherograms were processed using Peak Scanner Software v1.0 (Thermo Fisher Scientific). Protein structure prediction was performed using Phyre2.2 [145] and modelling of the WT and frameshift protein was performed using PyMOL [146]. The six lines and a Wm82 wildtype control were grown as outlined in *Plant material, microbial inoculation, and growth conditions* with 3 replications each. Plants were uprooted at 2 weeks and 4 weeks post emergence and roots were stained as outlined in *Root phenotyping*. 6 root segments per replicate were observed for AMF colonization, with ∼30 images taken per segment for quantification. AMF colonization percent was analyzed using AMFinder [147] outlined in D’Agostino, Raturi [37]. The experiment was repeated a second time with ∼5 replications each and the results remained statistically similar (Fig. 6g). Mean plots generated using ggplot2 were used for visualization and statistical analysis was performed using a pairwise Dunn’s test [148] within each experiment and time (2wpi & 4wpi) group, comparing each genotype to the corresponding Wm82 control.

---

**Fig. S1. Cluster Identification. (A)** Integrated UMAP projection of 33,410 high-quality nuclei across all five timepoints, resolved into 20 clusters. Each color represents a cluster as indicated in the legend. **(B)** Consensus UMAP plots of marker gene expression. Color intensity represents scaled average expression (yellow = low, black = high). Dotplot representing expression of marker genes in cluster 12. Dot size indicates the percentage of nuclei expressing each marker gene set; color intensity represents scaled average expression (blue = low, red = high).

**Fig. S2. Spatial metabolomics sectioning.** Images of root sections used for spatial metabolomics analysis. Boxes and labels denote specific samples.

**Fig. S3. Cortex defense GO term genes.** Dotplots of defense pathway gene expression across all cell-type sample groups. **(A)** regulation of immune response. **(B)** defense response to other organism genes. **(C)** DNA methylation. Dot size indicates the percentage of nuclei expressing each gene; color intensity represents scaled average expression (blue = low, red = high). Gene name legend is provided in table S7.

**Fig. S4. Cortex defense GO term genes continued.** Dotplots of defense pathway gene expression across all cell-type sample groups. **(A)** consensus dotplot of NBS-LRR classes. **(B)** induced systemic resistance. Dot size indicates the percentage of nuclei expressing each gene/group; color intensity represents scaled average expression (blue = low, red = high). Gene name legend is provided in table S7.

**Fig. S5. Gene Co-expression Network dotplot.** Consensus dotplot of GCN module expression across all cell-type sample groups. Dot size indicates the percentage of nuclei expressing each module; color intensity represents scaled average expression (blue = low, red = high). All genes in modules present in table S3.

**Fig. S6. Brown GCN network. (A)** visual representation of the brown GCN. Created in cytoscape. **(B)** GO BP term enrichment of network in **(A)**. Full gene list of all networks present in table S3. **(C)** Violin plot of brown network hub genes *GmNSL1, GmPAD4,* and *GmJAZ1*. Height of lines represent scaled expression level. Size of violin represents distribution of expression within the nuclei of the cell-type sample comparison groups; wider regions indicate a greater density of cells at that value.

**Fig. S7. Carbon metabolism pathways.** Dotplots of carbon pathway gene expression across all cell-type sample groups. **(A)** sucrose transport. **(B)** sucrose catabolism. Dot size indicates the percentage of nuclei expressing each gene; color intensity represents scaled average expression (blue = low, red = high). Gene name legend is provided in table S7.

**Fig. S8. Fatty acid and spatial metabolite profiles. (A)** Heatmap of top 50 expressed FA synthesis genes in the cortex. Color intensity represents scaled average expression (blue = low, red = high). All genes in modules present in table S7. **(B)** Consensus dotplot of selected saccharide and lipid compounds across all sample, colonized segment, and region comparison groups. Dot size indicates the percentage of pixels accumulating each compound; color intensity represents scaled average expression (blue = low, red = high).

**Fig. S9. Spatial metabolomics root segments of lipids.** Consensus root plots of C length lipids from spatial metabolomics. **(A)** long chain lipids (C13-C20), **(B)** very long chain lipids (C21-C34), **(B)** ultra long chain lipids (>C35).

**Fig. S10. Spatial metabolomics regions.** Plot of spatial metabolomics root segments with representative regions defined. Regions identified based on proximity to edge of root segments with EBImage package.

**Fig. S11. Expression dotplots of nitrogen transport genes.** Dotplots of nitrogen transport pathway gene expression across all cell-type sample groups. **(A)** nitrogen import. **(B)** nitrogen transport/assimilation. **(C)** ammonia assimilation. Dot size indicates the percentage of nuclei expressing each gene; color intensity represents scaled average expression (blue = low, red = high). Gene name legend is provided in table S7.

**Fig. S12. Expression dotplots of phosphorus transport genes.** Dotplots of phosphorus pathway gene expression across all cell-type sample groups. **(A)** phosphate starvation. **(B)** phosphorus transport. Dot size indicates the percentage of nuclei expressing each gene; color intensity represents scaled average expression (blue = low, red = high). Gene name legend is provided in table S7.

**Fig. S13. Analysis of Green GCN.** Visual representation of the green GCN. Created in cytoscape and GO BP term enrichment of network in. Full gene list of all networks present in table S3.

**Fig. S14. Expression dotplots of cortex specific genes:** Dotplots of cortex related gene expression across all colonized sample comparison groups. **(A)** select RdDM genes. **(B)** select HSC70 and hsp90.4 chaperone genes. **(C)** cell cycle and endoreduplication pathways. **(D)** endocytosis and exocytosis related pathways. Dot size indicates the percentage of nuclei expressing each gene/group; color intensity represents scaled average expression (blue = low, red = high). Gene name legend is provided in table S7.

**Fig. S15. Phenylpropanoid and coumarin dotplot.** Dotplot of phenylpropanoid and coumarin related gene expression across all cell-type sample groups. Genes with a red asterisk are genes from the red GCN module. The red box highlights the genes highly co-expressed with the red module genes. The purple asterisks represent the genes within significant QTLs from D’Agostino, Raturi [37]. Dot size indicates the percentage of nuclei expressing each gene/group; color intensity represents scaled average expression (blue = low, red = high). Gene name legend is provided in table S7.

**Fig. S16. Analysis of coumarin metabolites. (A)** Dotplot of coumarin compounds detected across all colonized root segment groups. **(B)** Root plot showing accumulation of dimethylfraxetin across all segments. Red asterisks represent colonized segments.

**Fig. S17. Mutation analysis using fragment analyzer.** Fragment analyzer results for all 6 frameshift soybean PIs and the Wm82 control. Fragment size of Wm82 is 410bp. Size for all frameshift lines is 408bp. X axis represents base pair length of fragment, y axis represents peak intensity. Ar = area under curve, ht = heigh of curve peak, sz = base pair size of fragment, dp = raw peak size from gel analysis.

**Table S1. QC metrics per sample.** Minimum, median, mean, and maximum UMIs and number of genes per nuclei in each sample.

**Table S2. Cell Type Markers for Soybean Roots.** Cell type markers used for identification in the current study, their respective cell type, and the study from which they were obtained. DOIs for each study provided.

**Table S3. GCN genes.** Genes from all GCN networks from Fig. S5. Present are the genes, module color, hub or member identity, Arabidopsis ortholog, gene name, and description, and soybean GO BP annotation.

**Table S4. AMF marker genes.** Colonized cell gene marker orthologs from Serrano, Bezrutczyk [13]. Contained are the gene names, soybean gene id, and DOI of original source.

**Table S5. Colonized cortex DEGs.** DEG list from Fig. 5. Contained are the stage, soybean gene id, Arabidopsis ortholog, gene name, and description, and soybean GO BP annotation.

**Table S6. Natural variant data.** Data used to generate Fig. 6g. Contained is the data point, sample, and percent colonization per data point. Data name legend included

**Table S7. Gene Fig. legend.** Legend of all gene names used in Fig. and their respective Fig. location

**Table S8. Primers for F6’H1 gene fragment amplification.** Primers used for gene fragment isolation. Included is the forward primer with M13 for fragment analyzer fluorescent labelling.

## Acknowledgements

G.B.P. is grateful to the United Soybean Board (Project # 25-209-S-D-2-B) and State of Texas’ Governor’s University Research (GURI) for the research funding. L.D’A is grateful to the Texas Tech University Graduate School.

## Authors Contribution

LDA: methodology, experimental design, formal analysis, data interpretation, and writing – original draft; K.G., genotyping and data interpretation; YL: methodology; LHE: experiment design and manuscript editing; GBP: conceptualization, formal analysis, funding acquisition, supervision, and writing – review & editing.

## Conflict of interest

G.B.P., L.D.A, and L.H.-E. are co-inventors on a patent application submitted in relation to the findings reported in this study.

## Declaration of generative AI in the writing process

During the preparation of this work, the author(s) used AI tools in order to synthesize the literature, correct the grammatical and language errors, and reconstruct some of the sentences to make them clear and easy to follow. After using this tool/service, the author(s) reviewed and edited the content as needed and take full responsibility for the content of the published article.

## Notes

### Summary of Updates

There was typo in the title and we have corrected it now.

