## Supplementary Figures for "A spatiotemporal single-cell atlas reveals coordinated immune, metabolic, and nutrient exchange programs and a coumarin-centered metabolic switch during soybean arbuscular mycorrhizal symbiosis"

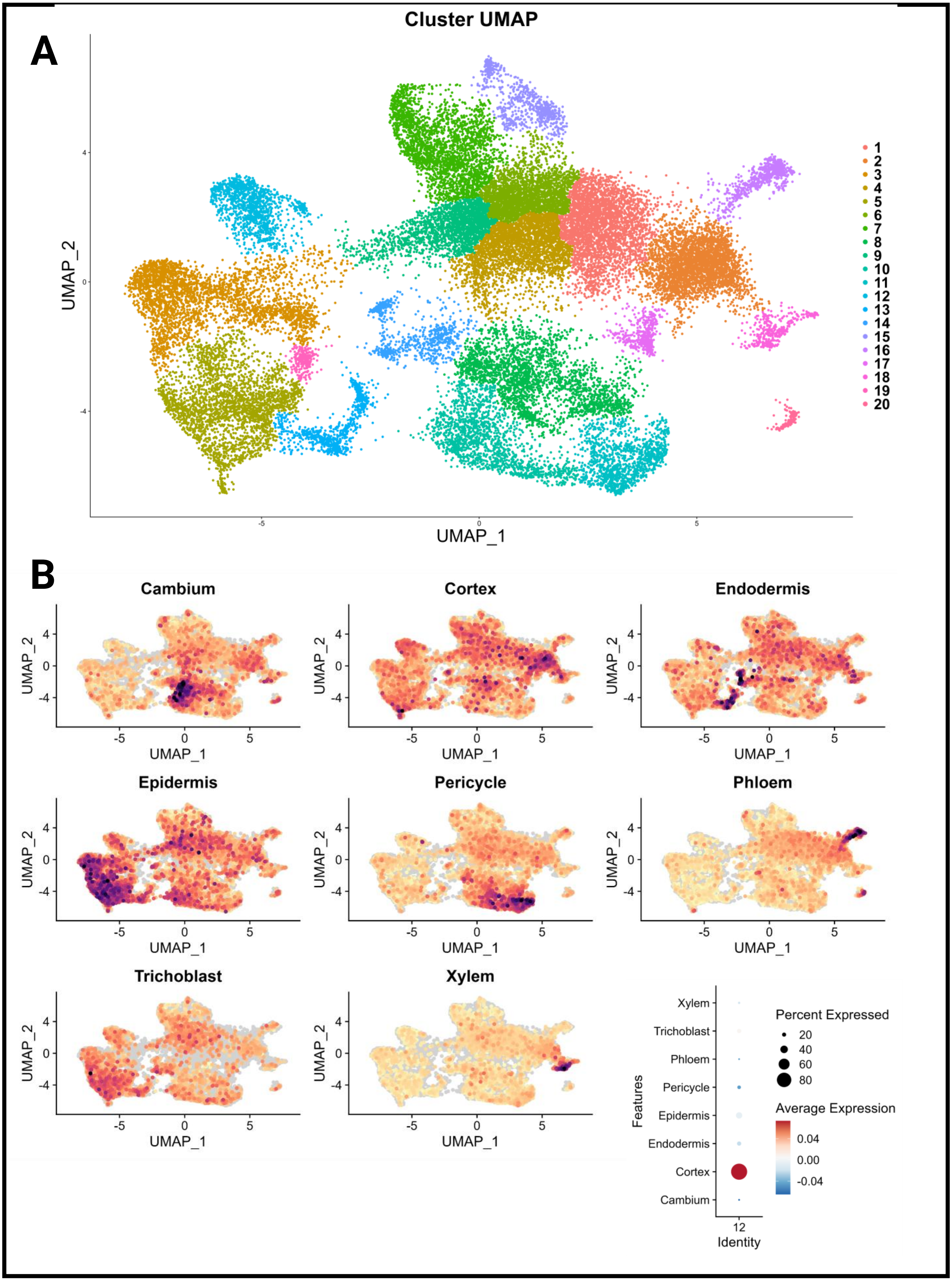


Figure S1: cluster identification

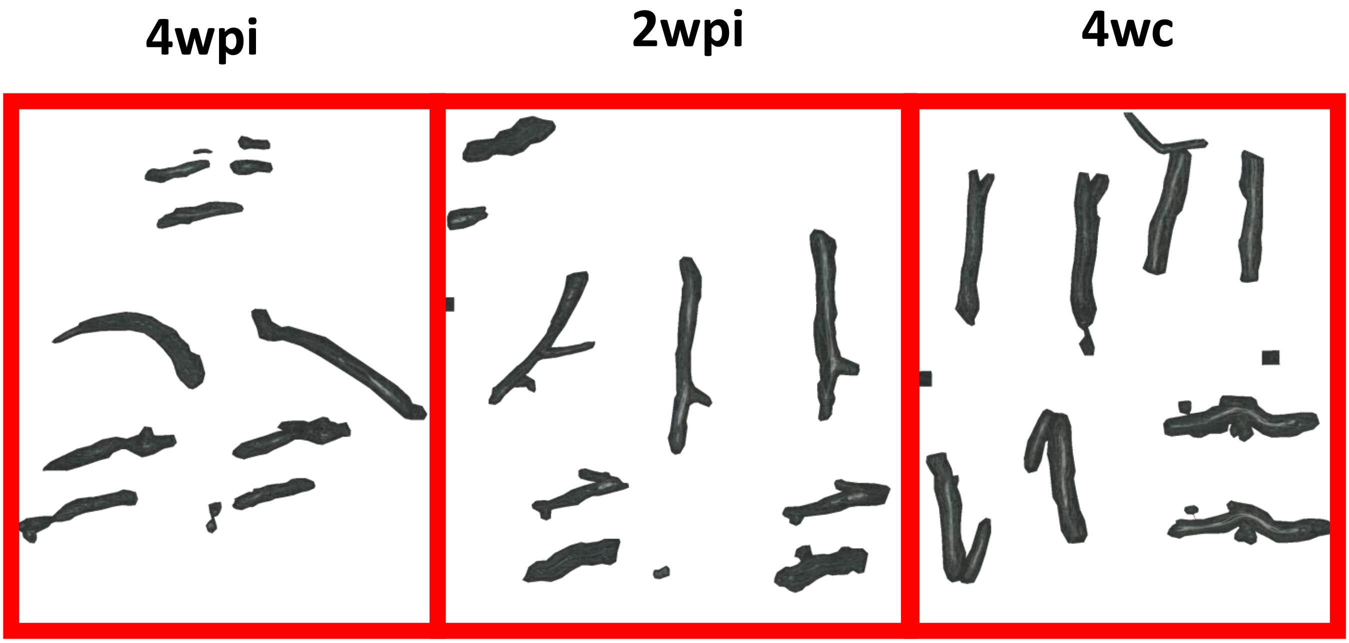


Figure S2: Spatial metabolomics root segments

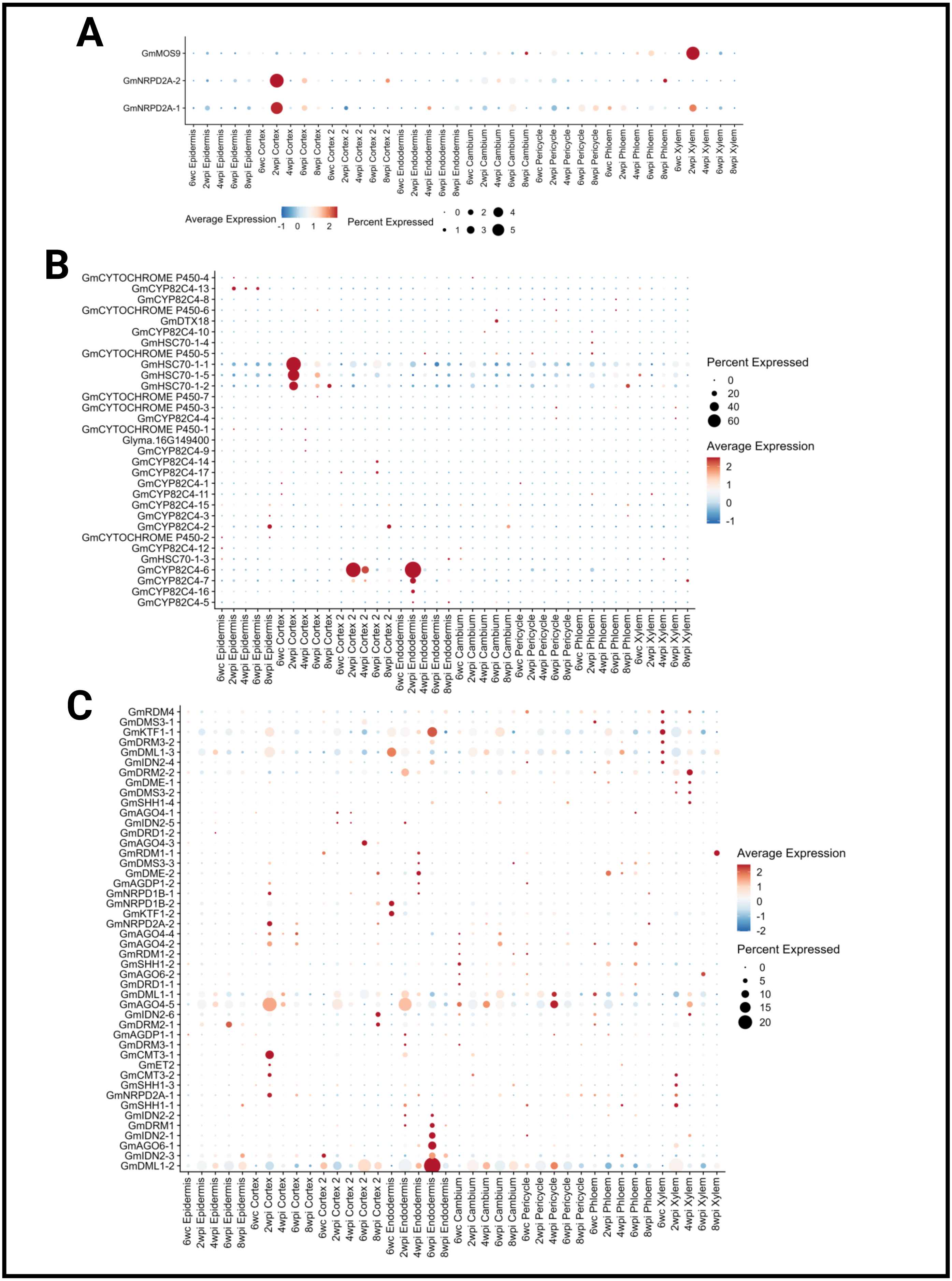


Figure S3: regulation of immune response, defense response to other organism genes, DNA methylation

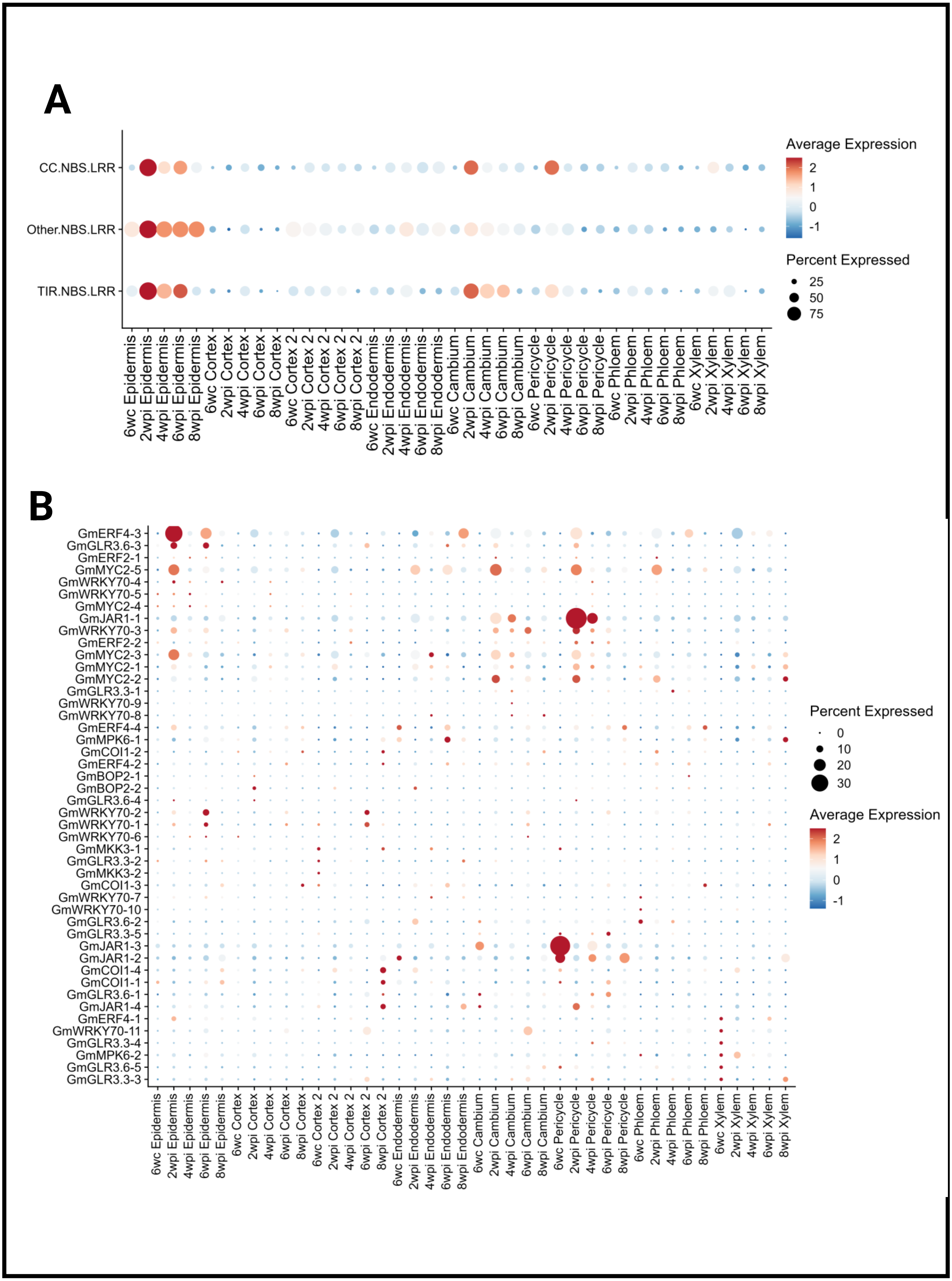


Figure S4: NBS-LRR + ISR

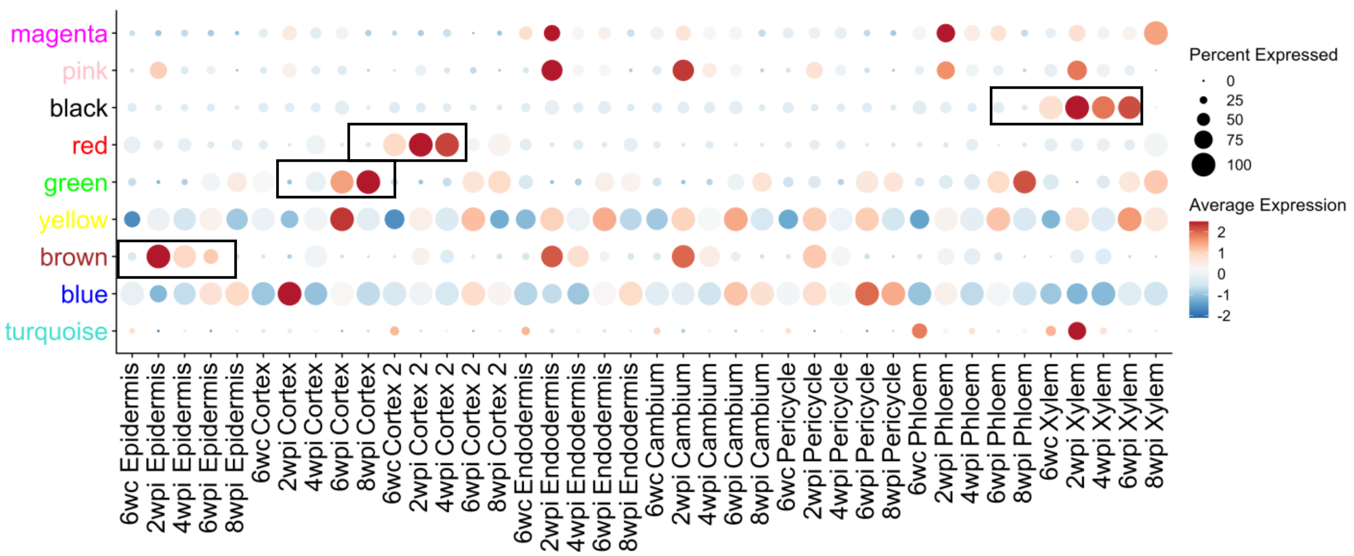


Figure S5: GCN dotplot

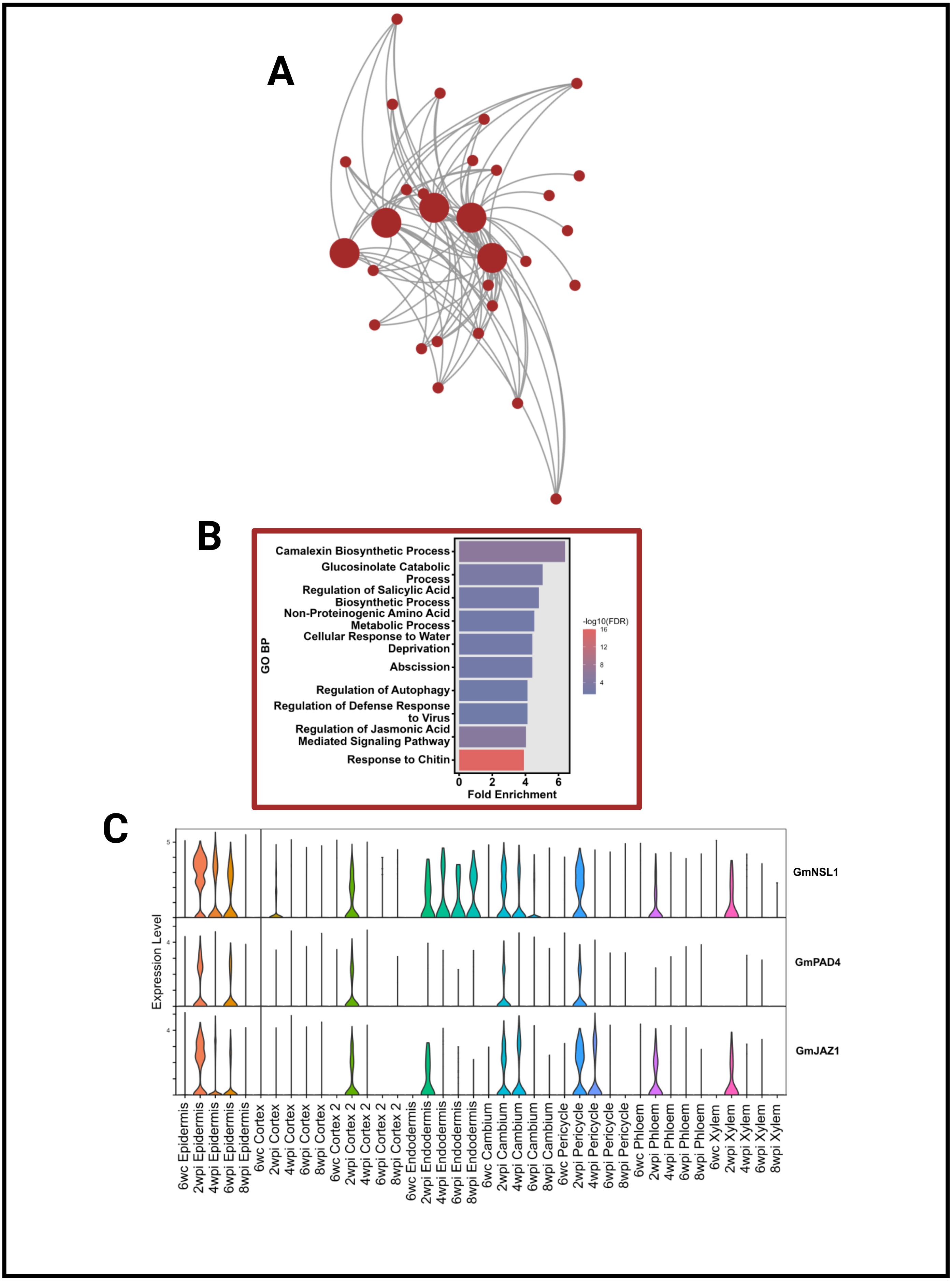


Figure S6: brown network

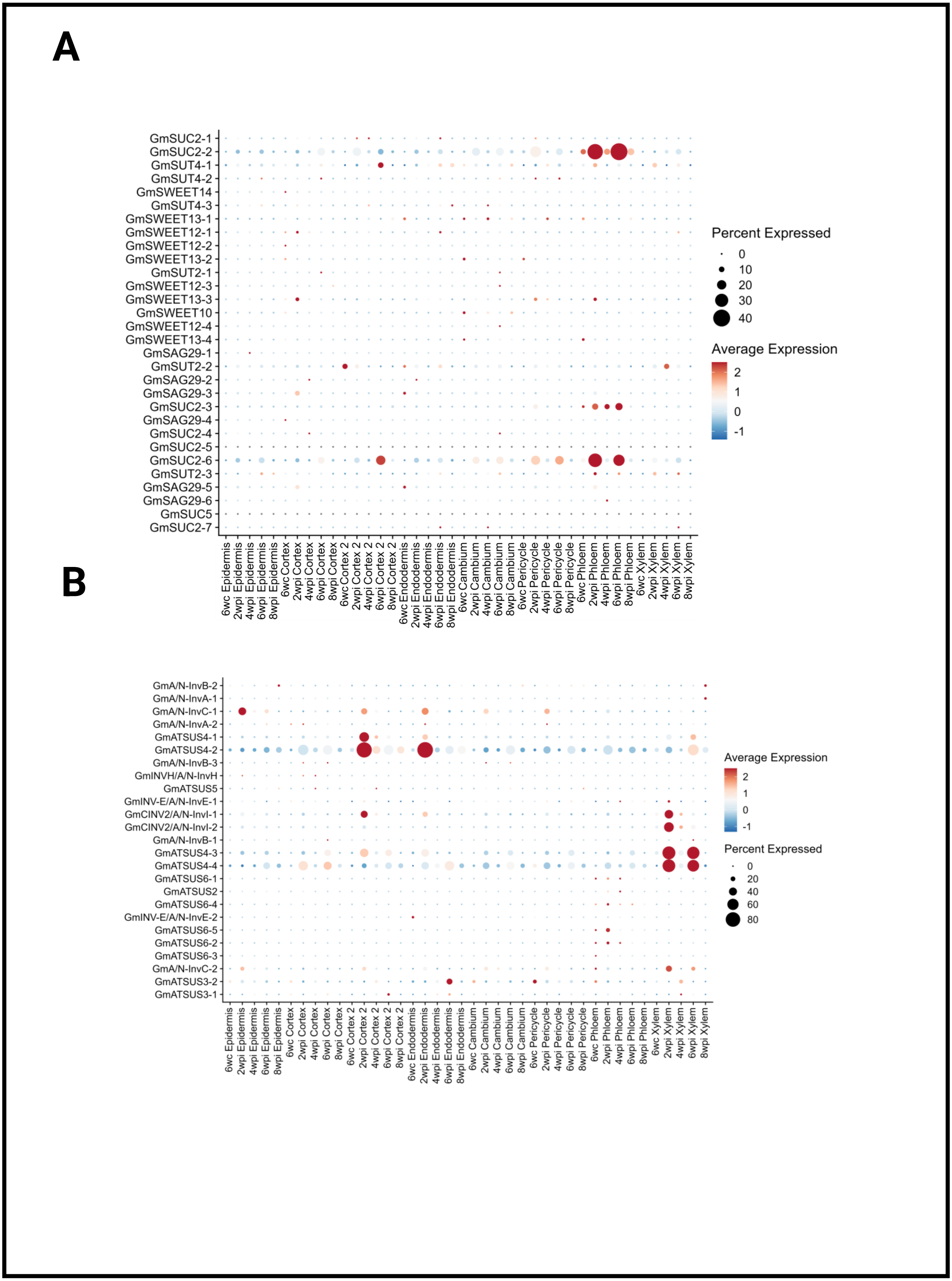


Figure S7: sucrose transport (a) + catabolism (b)

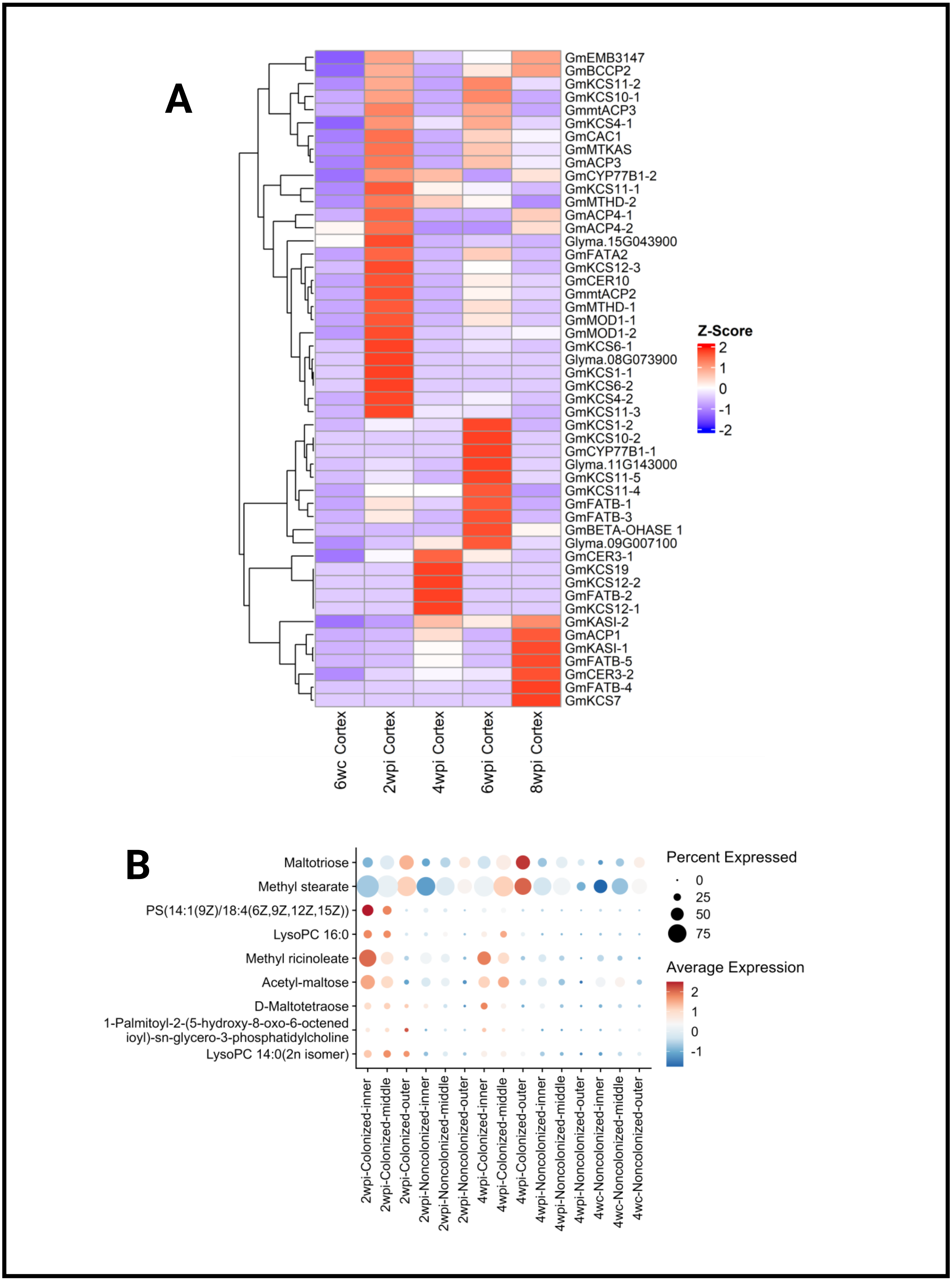


Figure S8: FA plots

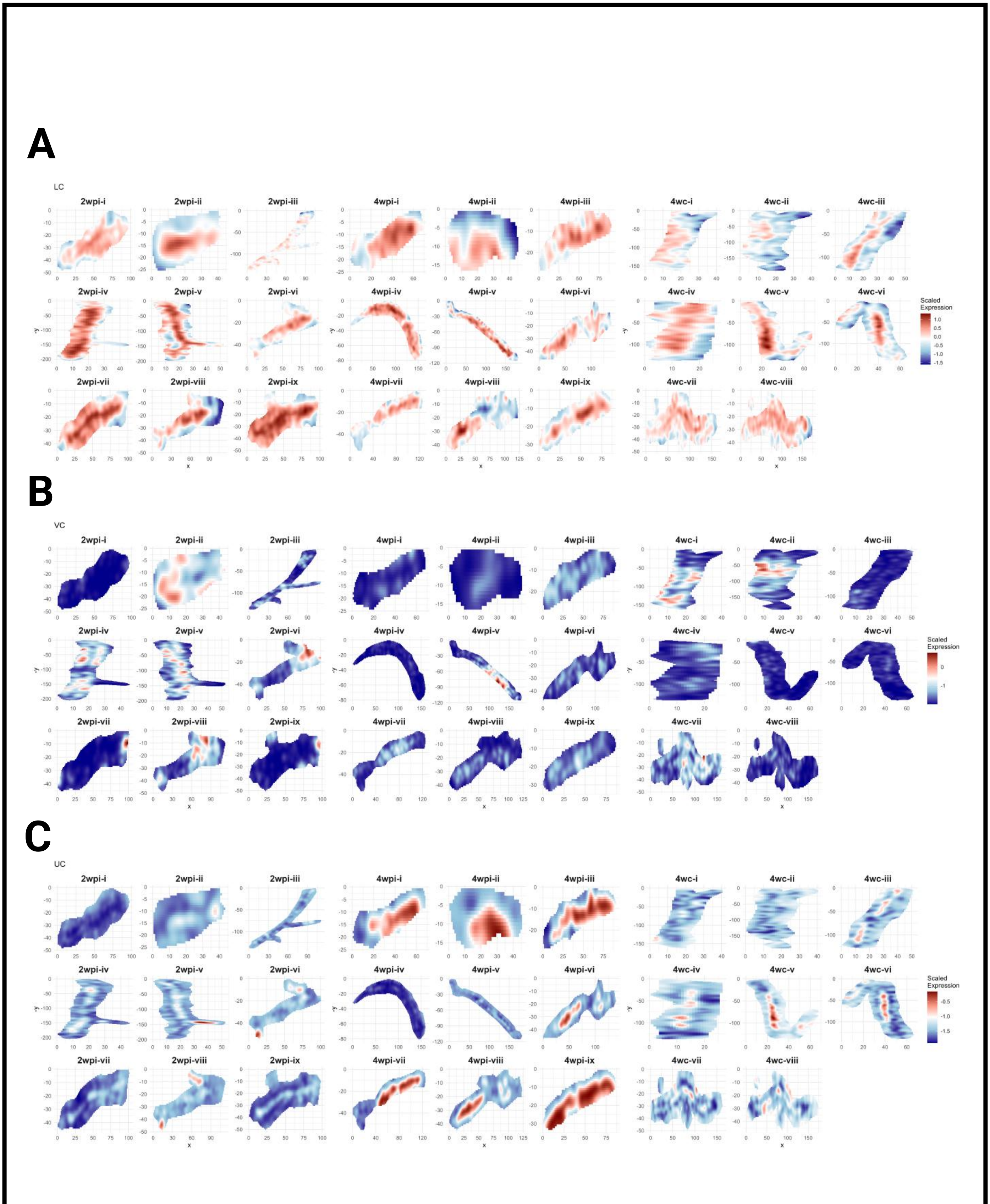


Figure S9: lipid root plots

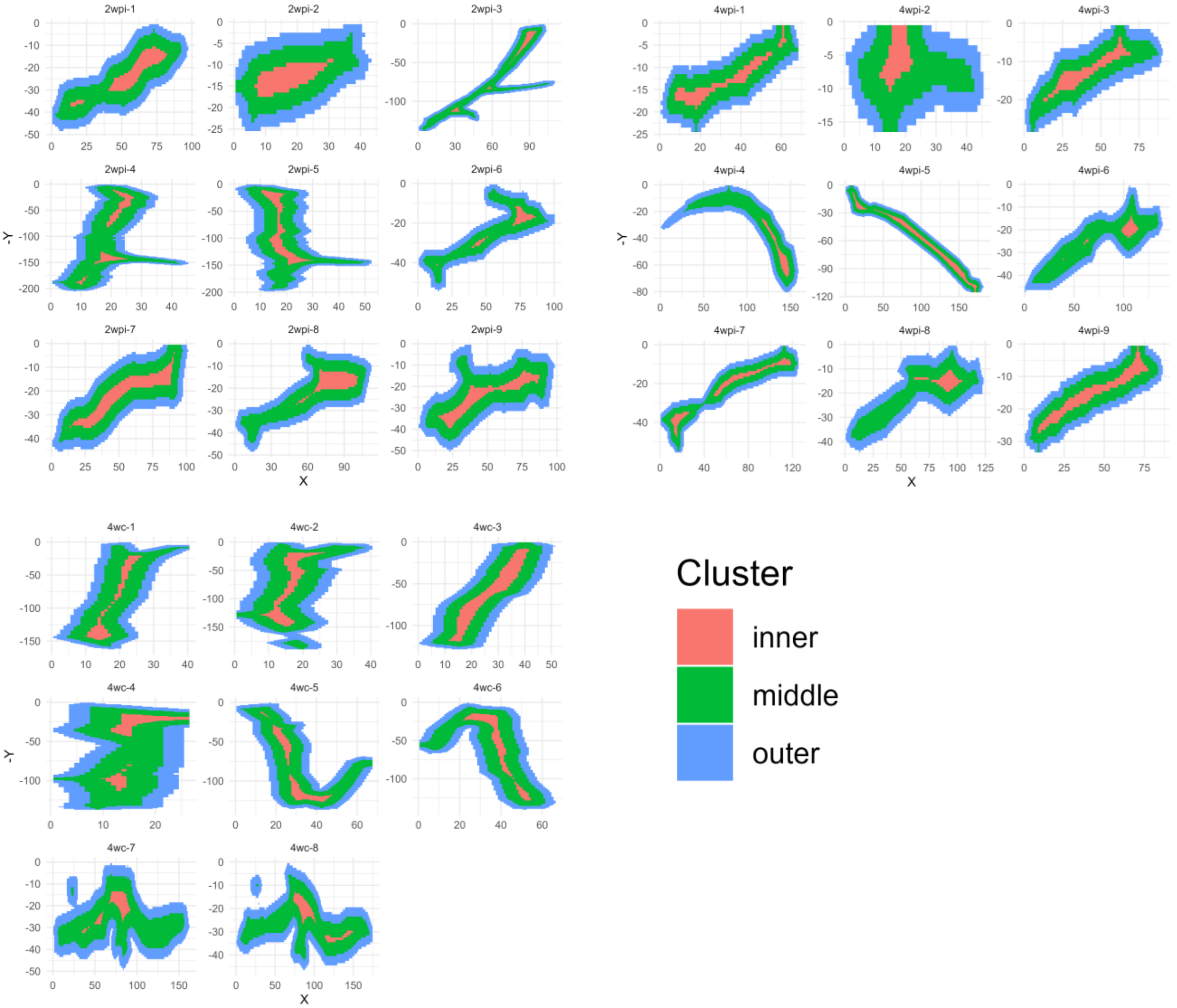


Figure S10: root segment regions

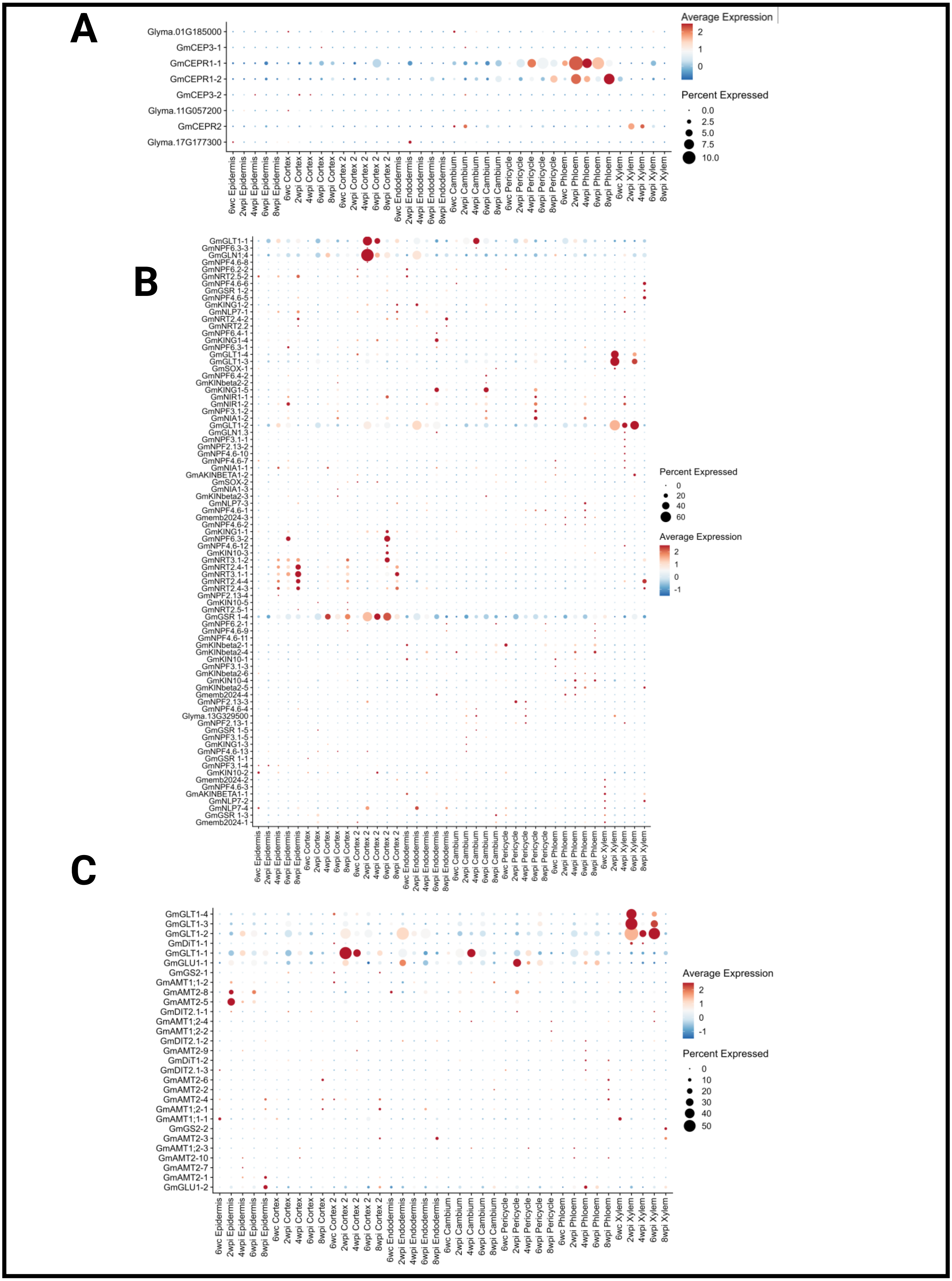


Figure S11: nitrogen import (a), nitrogen transport/assimilation (b), ammonia assimilation (c)

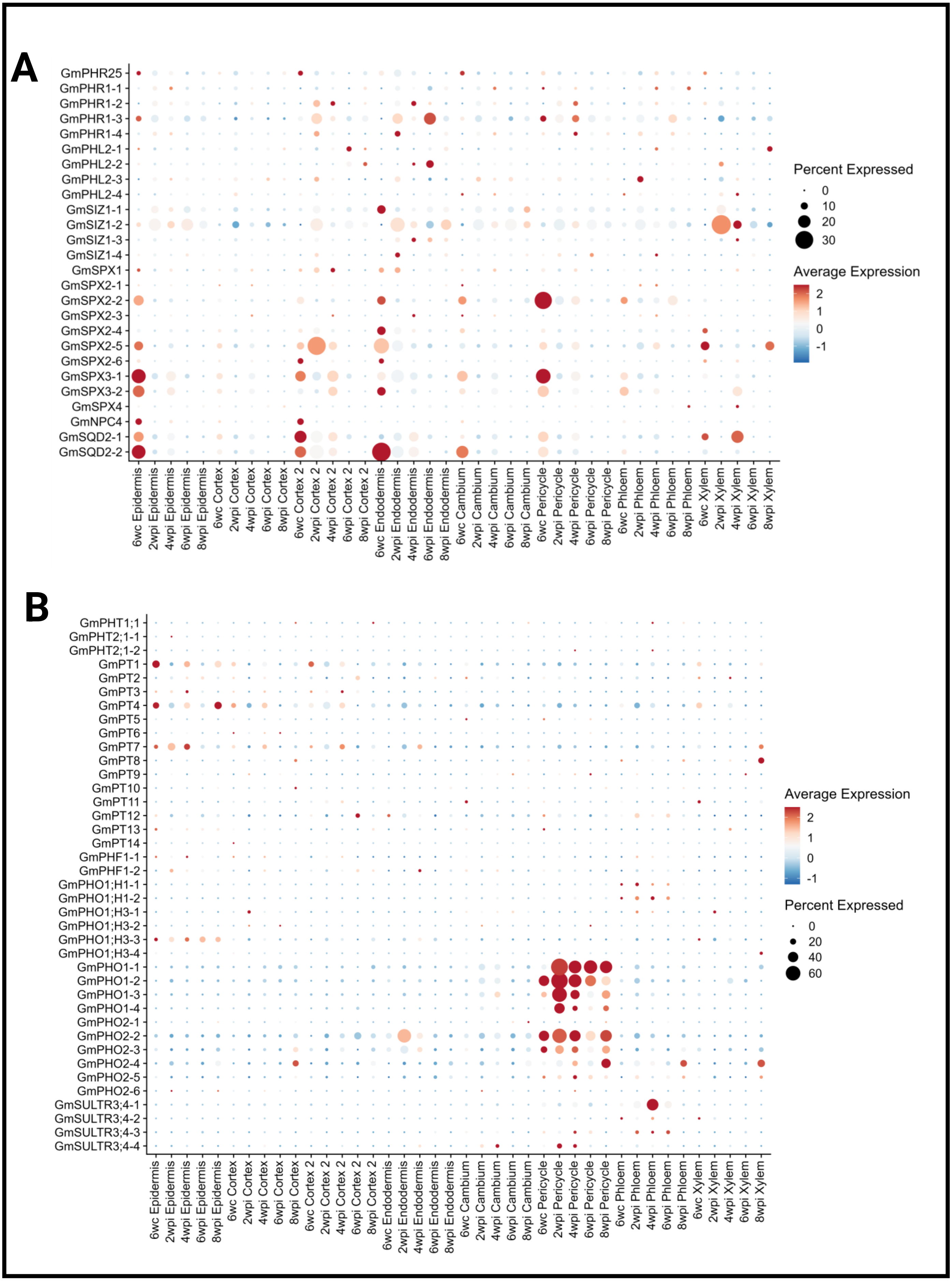


Figure S12: Phosphate starvation (a) and phosphate transport genes (b)

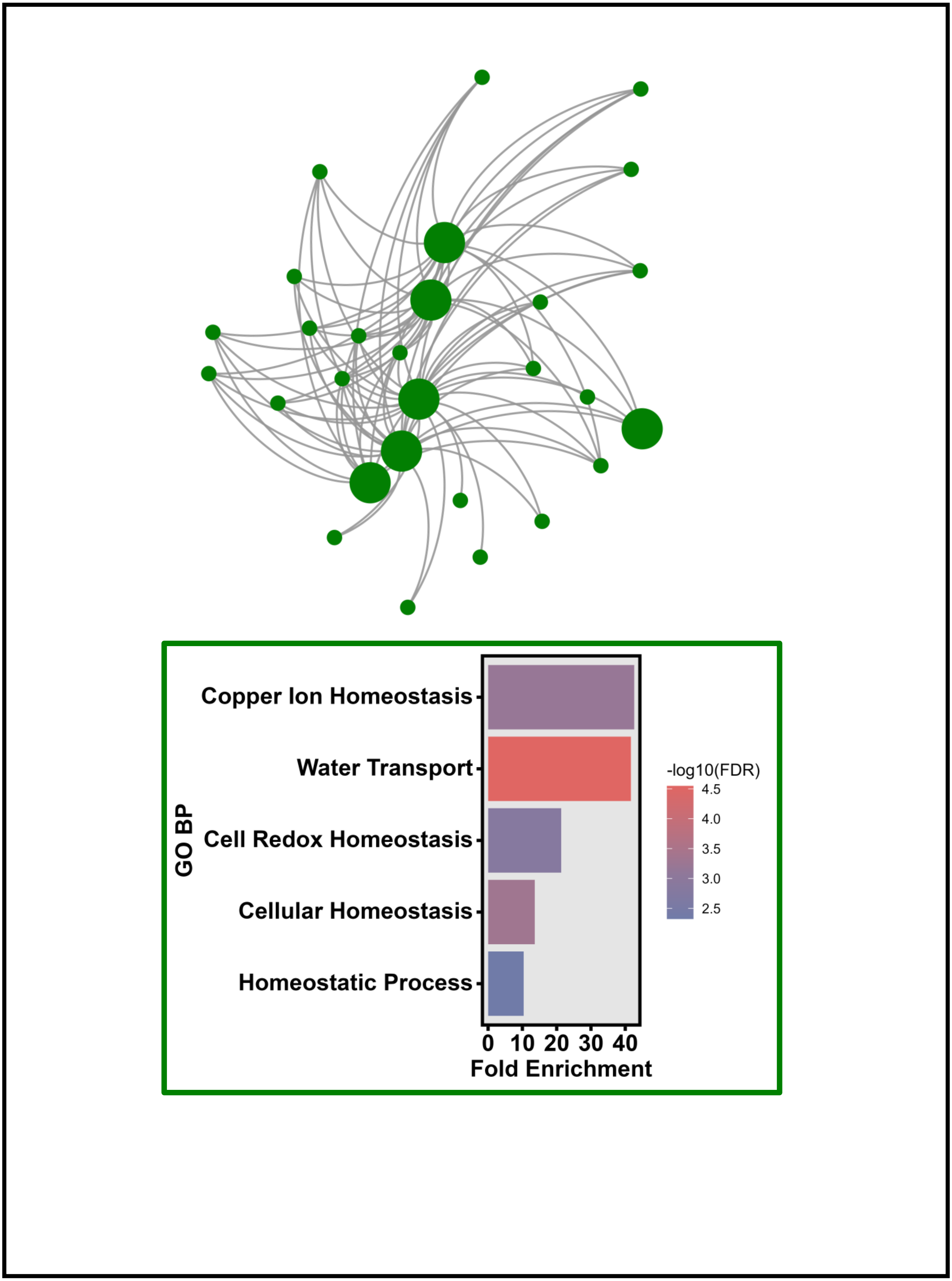


Figure S13: green network and GO term enrichment

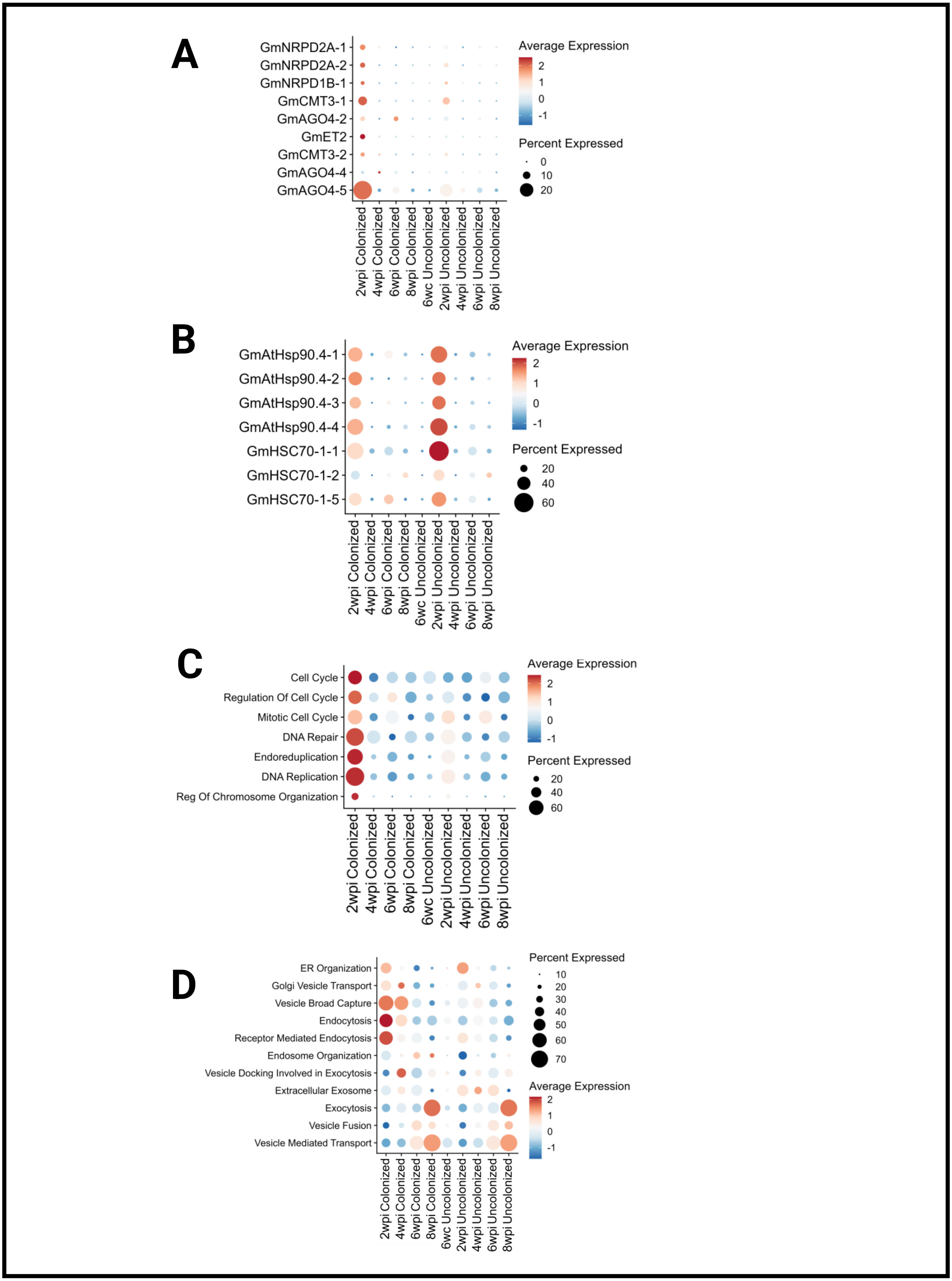


Figure S14: cortex explored fig 5

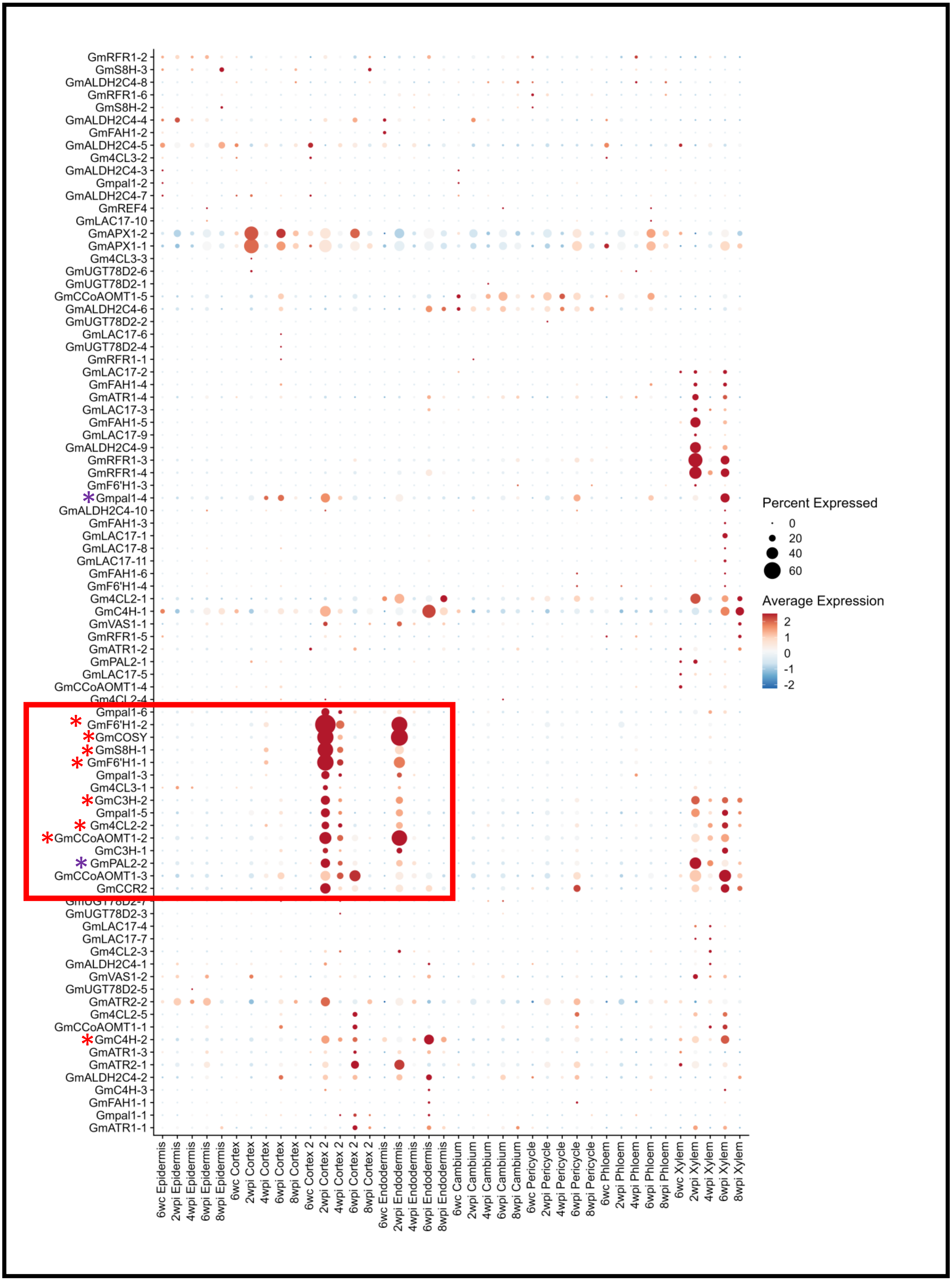


Figure S15: phenylpropanoid/coumarin dotplot. * = red network genes. * = GWAS genes

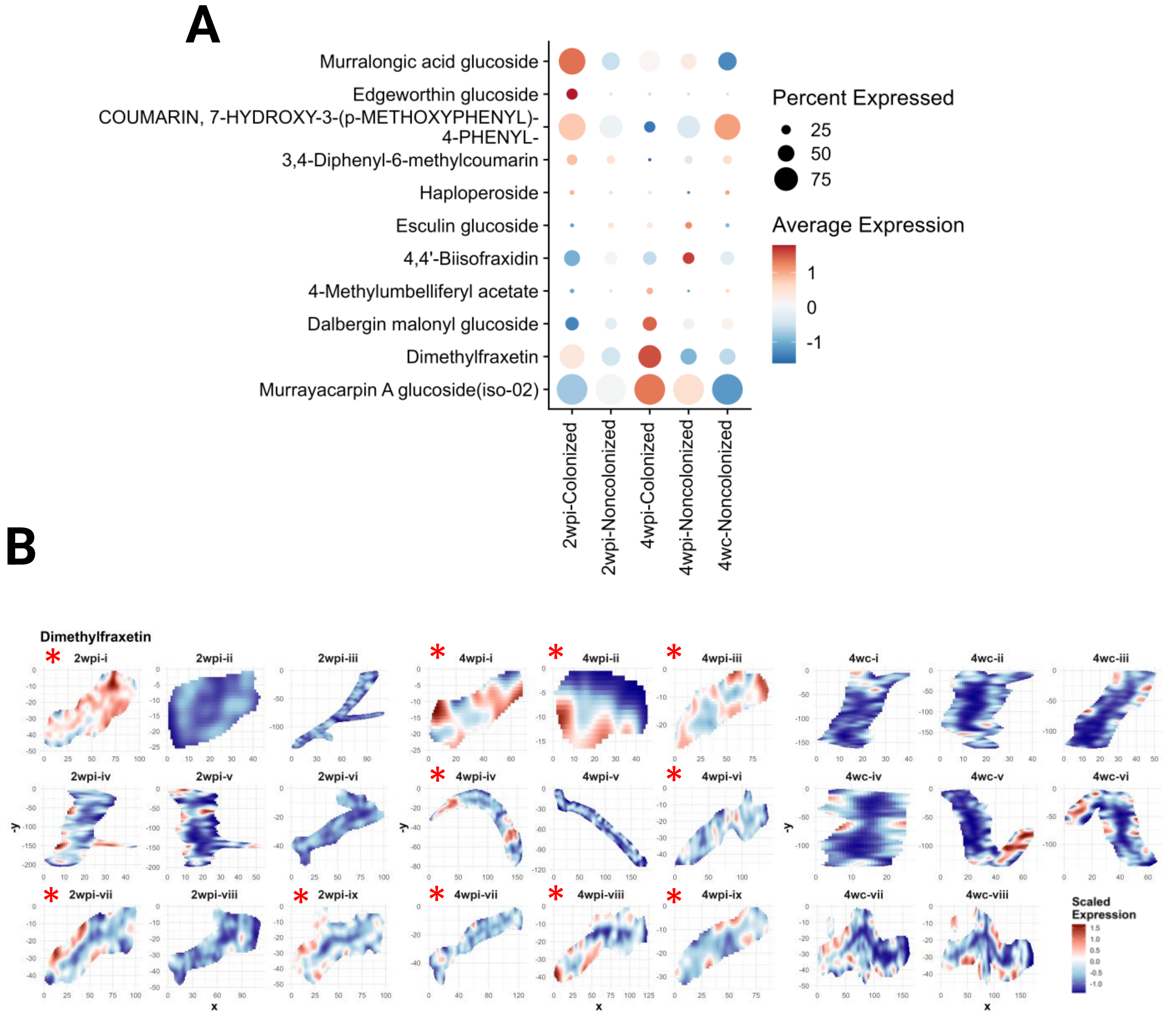


Figure S16: dimethylfraxetin all segments. * = colonized segments

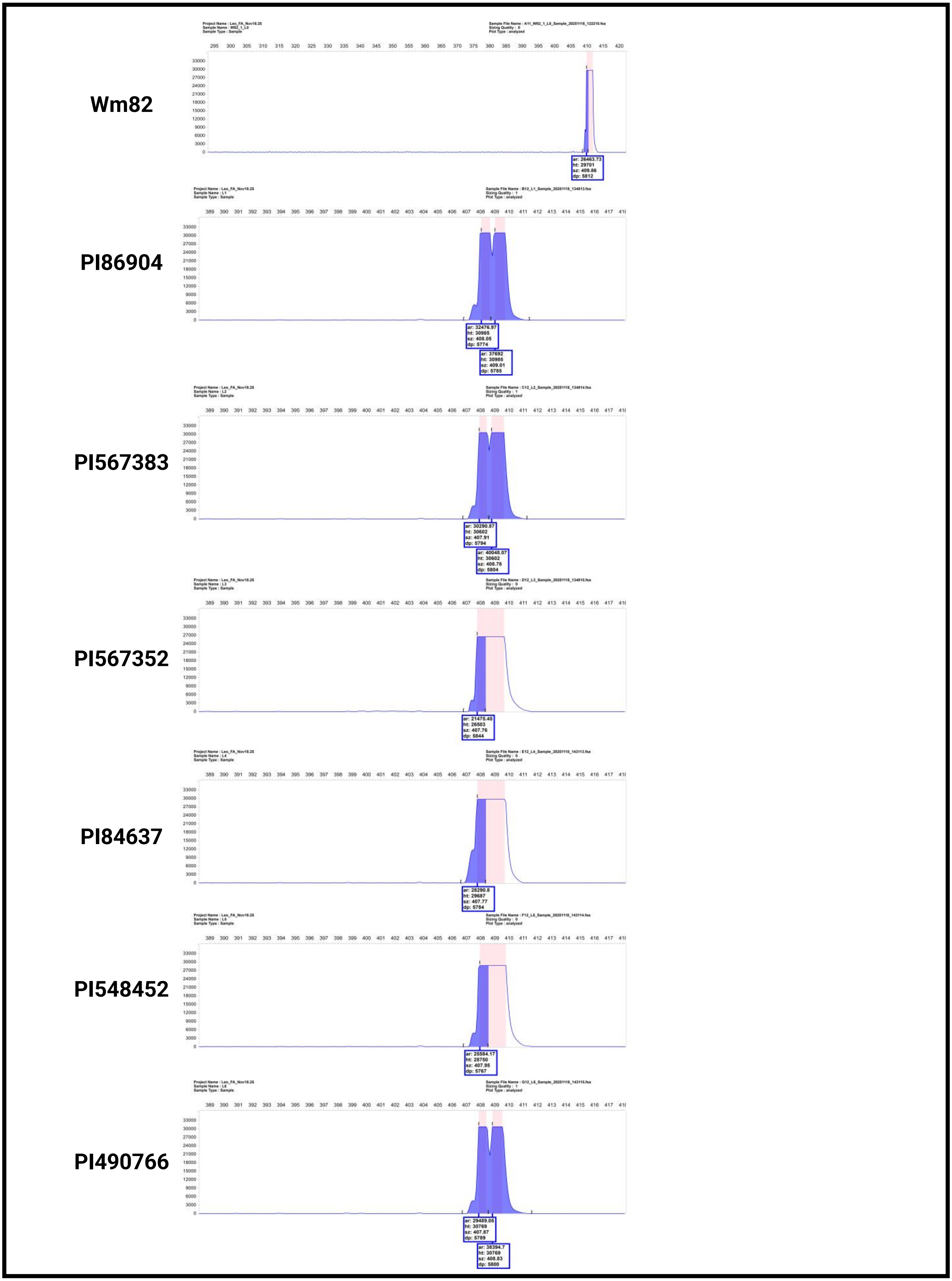


Figure S17: fragment analyzer analysis of frameshift in F6’H1 gene
